# The histone methyltransferase NSD3 contributes to sister chromatid cohesion and to cohesin loading at mitotic exit

**DOI:** 10.1101/2021.12.16.472961

**Authors:** Grégory Eot-Houllier, Laura Magnaghi-Jaulin, Gaëlle Bourgine, Fatima Smagulova, Régis Giet, Erwan Watrin, Christian Jaulin

## Abstract

Sister chromatid cohesion guarantees the correct transmission of chromosomes to daughter cells, and this multi-step process occurs throughout the cell cycle. Loading of the core cohesin complex onto chromatin takes place during mitotic exit, cohesion establishment happens during DNA replication, and the timely removal of cohesin occurs during mitosis. While cohesion establishment and mitotic cohesion dissolution have already been explored, the regulation of cohesin loading is not as well understood. Here, we report that the histone-lysine N-methyltransferase NSD3 is an essential factor in sister chromatid cohesion and mitotic progression, and that this occurs before and not after entry into mitosis. We establish that NSD3 interacts with the cohesin loader complex kollerin (NIPBL/MAU2), and that at mitotic exit it ensures proper levels of both MAU2 and cohesin itself on chromatin. In accordance with this newly described function in cohesin loading, we also show that NSD3 associates with chromatin in early anaphase, prior to the loading recruitment of MAU2 and RAD21, and that it then dissociates from chromatin when prophase begins. Going further, we also demonstrate that of the two NSD3 variants existing in somatic cells, it is the long isoform that is responsible for regulating kollerin and cohesin chromatin-loading, and that this isoform’s methyltransferase activity is required for efficient sister chromatid cohesion. Based on these observations, we propose that NSD3-dependent methylation contributes to sister chromatid cohesion by ensuring the proper recruitment of kollerin and thus loading of cohesin.

## Introduction

To ensure that replicated DNA is correctly transmitted to the daughter cells during mitosis, sister chromatids need to be held together until all of the chromosomes are correctly bi-oriented toward the opposite spindle poles. Cohesin is a multisubunit protein complex with a ring-like structure, and it can topologically link two chromatin fibres^1^. Throughout the cell cycle, the complex contributes to the dynamic regulation of genome organization, transcription, DNA damage repair, and sister chromatid cohesion. The core of the complex is made up of the three core subunits SMC1A, SMC3, and RAD21, and these are further bound by regulatory factors such as the Scc3 homologues SA-1 and SA-2^2,3^. Cohesin is loaded onto chromatin during exit from mitosis by the kollerin (NIPBL/MAU2) complex^4–9^ Nipped-B-like protein (NIPBL, also known as SCC2) is responsible for cohesin loading, while MAU2 sister chromatid cohesion factor (MAU2, also known as SCC4) facilitates the binding of that protein onto chromatin^10–13^. Before DNA replication, cohesin and chromatin association is dynamic, and it is actively removed by the WAPL cohesin release factor (WAPL) aided by scaffolding proteins PDS5A and PDS5B^14–19^ Afterwards, during DNA replication, sister chromatid cohesion is established when a fraction of cohesin stably associates with chromatin through the acetylation of SMC3 by the ESCO1 and ESCO2 acetyltransferases^20–25^. This process is accompanied by a specific cohesin loading mechanism wherein kollerin binds to a phosphorylated form of the MCM2-7 pre-replication complex^26^. Acetylated SMC3 allows for the subsequent binding of sororin (also known as cell division cycle associated 5 or CDCA5) to cohesin complexes, where it antagonizes WAPL anti-cohesive activity until mitosis begins^19,27,28^. Upon mitotic entry in human cells, the mitotic kinases CDK1, PLK1, and Aurora B phosphorylate sororin and the cohesin components. This makes cohesin sensitive to the activity of WAPL, and in a process known as the prophase pathway^29,30^, the cohesin molecules disassociate from chromosomes arms. During this process, sister chromatid cohesion is protected from WAPL-mediated dissociation at the centromere by the proteins shugoshin 1 (SGO1) and the histone H3 associated protein kinase (HASPIN). SGO1 competes with WAPL for binding to the cohesin ring, while the SGO1 partner PP2A, the holoenzyme protein phosphatase 2A, is thought to counterbalance any phosphorylation of cohesin or sororin^31,32^. At the same time, HASPIN binds to PDS5 and prevents cohesin from interacting with WAPL^33,34^. Once all kinetochores are properly attached to microtubules during metaphase, the spindle assembly checkpoint (SAC) is turned off, and this leads to the activation of the endoproteinase separase that cleaves the RAD21 cohesin subunit, thereby causing the cohesin ring to open and the two sets of chromosomes to segregate^35^. Of note, in the case of prolonged mitotic arrest with centromeres under spindle tension, chromosomes are subjected to an unscheduled dissociation of the sister chromatids, a process referred to as cohesion fatigue^36,37^

In a previous work, we reported the presence of dimethylated H3K4 at the centromere in correlation with the prevention of premature sister chromatids separation (PSCS) during mitosis^38^. In determining whether histone-lysine N-methyltransferases are involved in sister chromatid cohesion, we identified the possible contribution of the H3K36-specific methyltransferase NSD3 to this regulation. NSD3 belongs to the nuclear receptor-binding SET domain protein subfamily, a group with three members: NSD1; NSD2, also known as WHSC1 (Wolf-Hirschhorn syndrome candidate 1); and NSD3, also known as WHSC1L1 (WHSC1-like 1). NSD methyltransferases act as oncoproteins in different types of cancers^39,40^, and they are considered to be specific for mono- or di-methylation of H3K3 6^41–45^, an epigenetic mark involved in gene transcription, RNA alternative splicing, and DNA replication, repair, and methylation. However, the contribution of NSD3 to this methylation seems to be lower than that of NSD2^46^. In addition to their SET domains, NSD family members are characterised by the presence of seven domains that bind to modified histones. Five of these are plant homeodomain (PHD) domains which recognize specific DNA sequences together with histone post-translational modifications (methylated lysine or arginine, acetylated lysine), and the other two are proline- and tryptophan-rich (PWWP) domains which bind methylated lysines^47–52^.

In somatic cells, NSD3 messenger RNA contains 24 exons and induces the expression of two protein isoforms, NSD3-long (NSD3-L) and NSD3-short (NSD3-s)^47,48,53^, while a third isoform WHISTLE (found in testes) is specifically expressed from a downstream promoter^48^. The long isoform NSD3-L contains 1437 amino acids (aa) and the SET methyltransferase domain is located in its carboxy-terminus. The short isoform NSD3-s (647 aa) is translated from an alternatively spliced messenger and since it lacks the SET domain it also lacks methyltransferase activity. NSD3-s also differs NSD3-L in its last 620-647 aa, and in having a single PWWP domain. That single NSD3-s PWWP domain is required for the function of BRD4-NSD3s-CHD8 complex to sustain leukemia cell proliferation^54^

Here, we report that inactivation of the methyltransferase SET-family member NSD3 results in PSCS during early mitosis. We show that NSD3 is required for cohesion in G2 phase before mitotic entry, and that it interacts with kollerin and improves cohesin loading at mitotic exit. We also describe that NSD3 loads onto chromatin in early anaphase, prior to kollerin recruitment. Finally, we also demonstrate that the protein’s role in cohesin loading is mediated by the NSD3-L long isoform, and that an active SET methyltransferase domain is required.

## Results

### NSD3 is required for sister chromatid cohesion

In order to identify new methylation effectors involved in regulating sister chromatid cohesion, we performed an inactivation screen with RNA interference (RNAi) to identify defective mitotic cohesion in 14 human SET domain-containing methyltransferases. For each methyltransferase, three different siRNAs were transfected in HeLa Kyoto cells. We prepared mitotic chromosome spreads, and analysed them for mitotic cohesion defects (Figure 1A). NSD3 was the only methyltransferase tested that produced a significantly increased proportion of cells displaying separated sister chromatids during prometaphase with all three tested siRNA (Figure 1B, Figure S1). The siRNAs used for the screen, 1, 2, and 3 targeted NSD3 exons 5, 6, and 2, respectively. As expected, since the short isoform of NSD3 results from an alternative splicing event between exons 9 and 10, these siRNAs decreased expression of both isoforms (Figure 1C). This screen therefore revealed that depleting NSD3 causes defective sister chromatid cohesion, and that this is detectable in mitosis.

**Figure 1:**
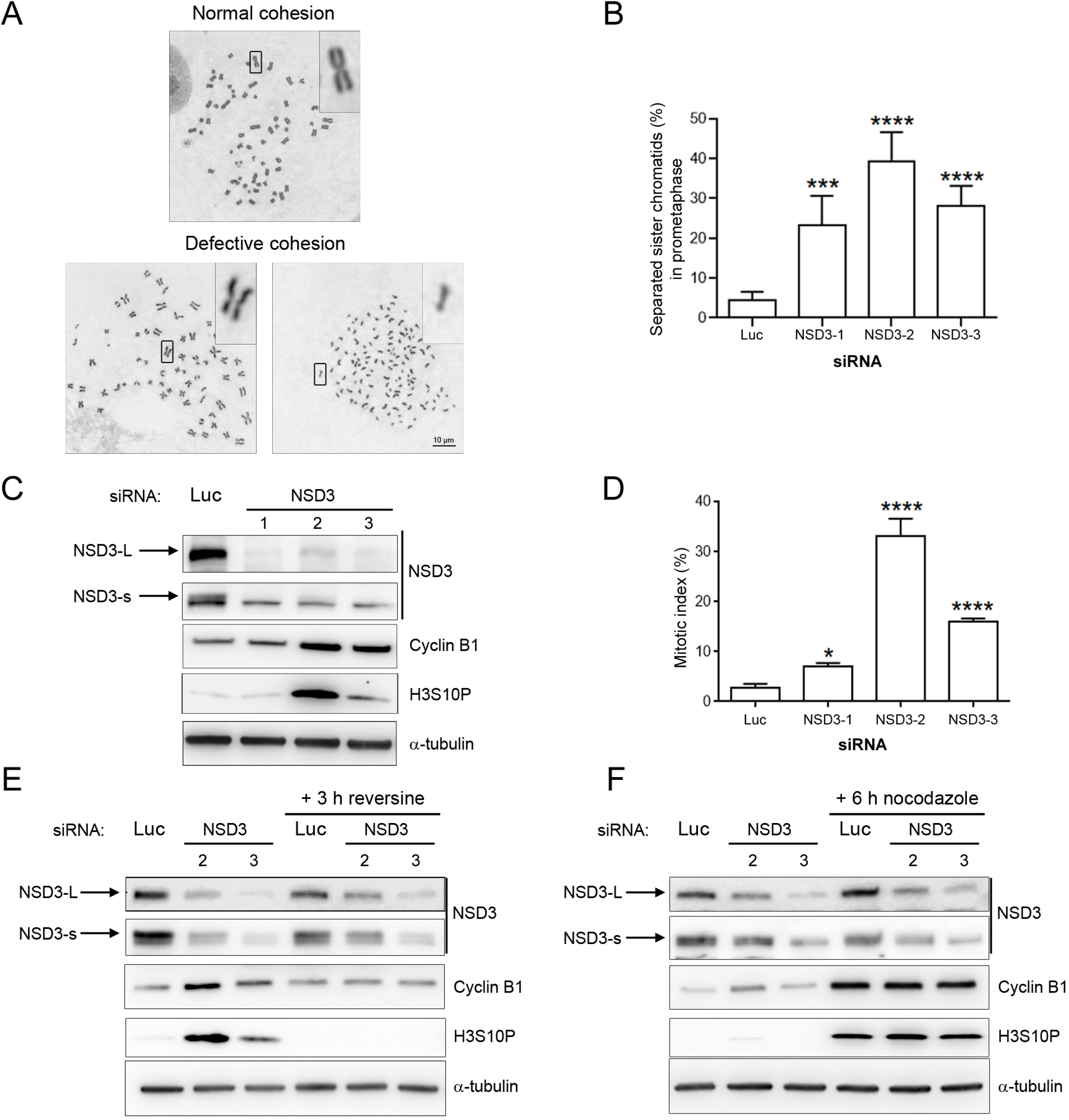
NSD3 depletion induces premature sister chromatid separation and SAC-dependent mitotic arrest. (A) Representative images of chromosome spreads of magnified chromosomes from HeLa cells transfected for 72 h with a luciferase or various SET-domain methyltransferase siRNAs. (B) Relative amounts of cells containing cohesion defects induced after NSD3 depletion (*n* =5, ≥ 300 prometaphase cells were counted in each condition). Error bars represent SD of the mean. ***: *P* ≤ 0.001; ****: *P* ≤ 0.0001. (C) Western blot (WB) analysis of NSD3 depletion after 72 h transfection with three different siRNAs. Long (NSD3-L) and short (NSD3-s) isoforms are indicated. Note the presence of a non-specific band just below the short isoform band. (D) Accumulation of mitotic cells in absence of NSD3 (*n* =3, ≥ 500 cells counted for each siRNA tested). Error bars represent the SD of the mean. *: *P* ≤ 0.05; ****: *P* ≤ 0.0001. (E) Effect of SAC inhibition on the mitotic cell accumulation induced by NSD3 depletion. As (C), except that cells were treated for 3 h with reversine before being harvested. (F) Effect of NSD3 depletion on the maintenance of SAC activity. Same as above, this time with 6 h of treatment with nocodazole.

Since NSD3 is involved in gene expression regulation, the cohesion defects that we observed could be caused by an altered expression of cohesin components or regulators. To test for this, we did immunoblot experiments to analyse the total amounts of the core cohesin subunits RAD21, SA2, and SMC1A, the cohesin loader MAU2, and the cohesin regulators sororin, ESCO2, and WAPL. We observed that NSD3 depletion did not alter the expression levels of any of these cohesin subunits and partners (Figure S2). Moreover, these experiments also indicated that global di-methylation levels of H3K36 remain unaffected by NSD3 inactivation (Figure S2), indicating that NSD3’s contribution to sister chromatid cohesion is independent of both the global disruption of H3K36 di-methylation and the altered expression of cohesin components and regulators.

We went on to examine the proportions of mitotic cells in the population, and saw that, depending on the siRNA used, NSD3 depletion causes a significant increase in the mitotic index (Figure 1D). An increase in the proportion of mitotic cells which exhibit separated sister chromatids, such as the one observed with NSD3 inactivation, can arise either because of an accumulation of prometaphase cells with defective mitotic cohesion, or because the cells have prematurely entered into anaphase. In order to discriminate between these two possibilities, we analysed the expression of two mitotic markers whose levels decrease when cells enter anaphase. The first was cyclin B1, whose levels increase starting at late S-phase, culminate during the first steps of mitosis, then rapidly degrade after anaphase onset. The second marker was phosphorylation of H3S10, a process which begins when the cells enter prophase and which is then rapidly erased after the transition from metaphase to anaphase. Following depletion with the two siRNAs that induced the strongest mitotic index increase, we observed an accumulation of both of these markers, indicating that cells were blocked in a prometaphase-like state (Figure 1C). To confirm that this arrest resulted from SAC activity, we treated NSD3-depleted cells with the SAC inhibitor reversine for 3 h (Figure 1E). In this case, the levels of both mitotic markers were similar to those in cells treated with the control siRNA. Moreover, we also treated cells with the microtubule-depolymerising agent nocodazole for 6 h, artificially ensuring that the SAC remained active, and there was again no difference after NSD3 depletion (Figure 1F). It is thus clear that NSD3 depletion does not affect SAC activity. Taken together, our results strongly suggest that cohesion defects induced by NSD3 depletion cause a SAC-dependent arrest during prometaphase in HeLa cells.

### NSD3 is involved in regulating the loading of cohesin

We next looked at which stage of the cohesion cycle is affected by NSD3 depletion. Because NSD3 inactivation induces PSCS, we excluded the idea that it contributes to the cohesion-releasing prophase pathway, as its depletion would lead to the opposite phenotype (a bivalent chromosome with cohesive arms rather than an X-shape). We therefore looked for the presence and localization of the centromeric cohesion protector SGO1. After NSD3 depletion, we detected as much SGO1 at the centromeres of separated sister chromatids as we saw in control cells having attached sister chromatids (Figure 2A). However, two pools of SGO1 localized at the inner and outer centromeres have been described^55^. The inner pool is required to protect centromeric cohesion, but the function of the outer pool remains elusive^32,56^. In the absence of NSD3, sister chromatid separation prevented us from determining whether the centromeric SGO1 we observed corresponded to one or both of these pools. Protection of centromeric sister chromatids also depends on the kinase HASPIN, via WAPL phosphorylation and HASPIN’s interactions with PDS5 and WAPL^33,56,57^. Whether the molecular mechanism involves SGO1 or HASPIN, the protection of centromeric cohesion occurs by counteracting WAPL. Mitotic cohesion fatigue is also sensitive to the presence of WAPL^36^. We therefore explored the effects of WAPL depletion in the absence of NSD3. To verify the functional efficiency of such a depletion, we confirmed that co-depletion of HeLa cells with both WAPL and SGO1 siRNA prevented PSCS, while cells depleted for SGO1 alone caused it (Figure S3A). As an additional control, we also showed that the co-depletion of WAPL and RAD21 fails to rescue PSCS which had been induced by the lack of the core cohesin components (Figure S3B). Using the same set-up, we were able to show that WAPL depletion does not rescue PSCS induced by NSD3 depletion (Figure 2B). Together, these results indicate that NSD3 is not involved in maintaining sister chromatid cohesion during mitosis.

**Figure 2:**
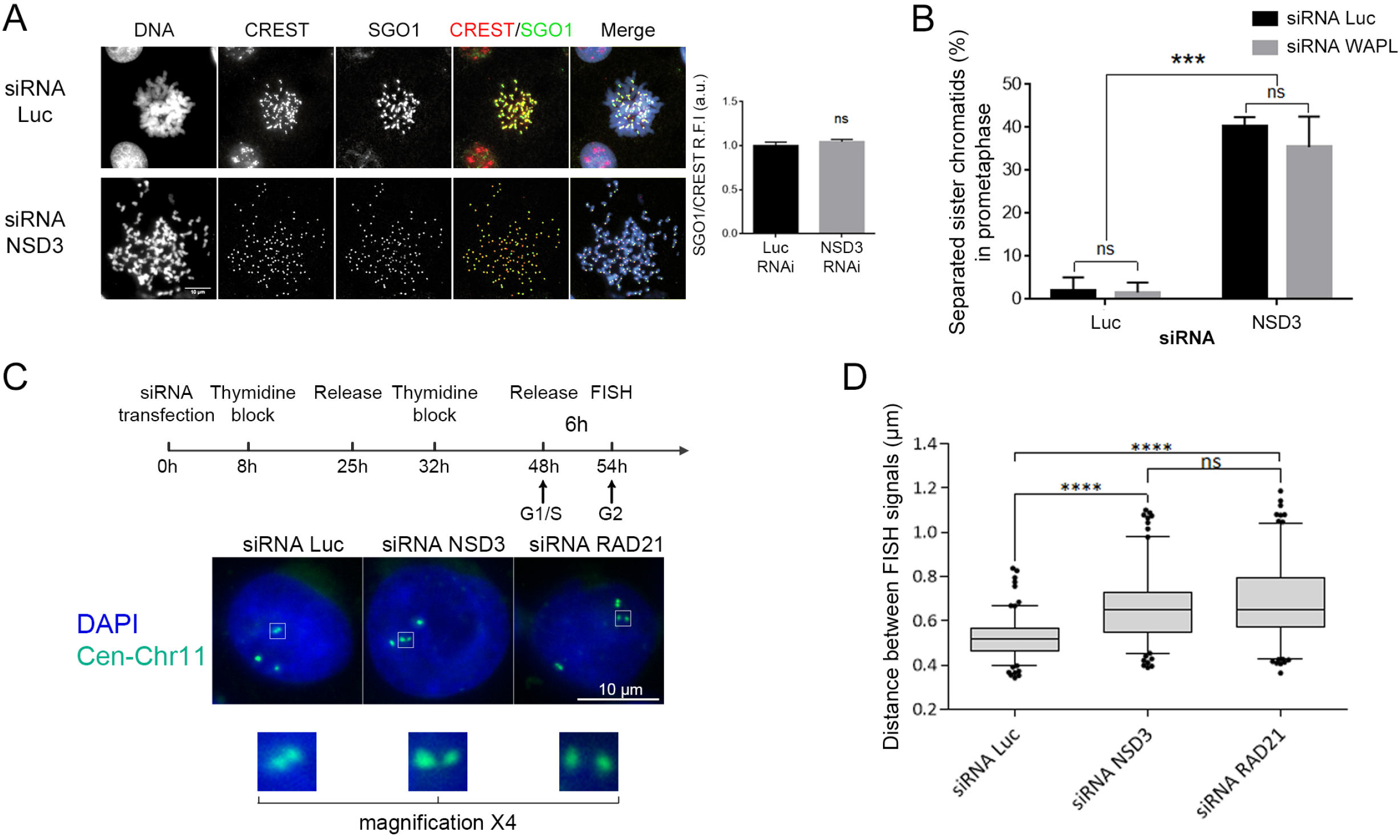
The methyltransferase NSD3 contributes to cohesion maintenance before mitotic entry. (A) *Left*, Immunofluorescence with antibodies against the centromeric cohesion protector SGO1 after 72 h of treatment with luciferase (Luc) or NSD3 siRNA. CREST serum was used to label the positions of the centromeres. *Right*, SGO1 fluorescence intensity relative to that of CREST (R.F.I) shown in arbitrary units (a.u.) as the mean ± SEM of a representative experiment (*n* ≥ 50). ns: not significant. (B) In prometaphase, the percentage of separated sister chromatids in cells depleted for NSD3, WAPL, or both (*n* =2, ≥ 150 prometaphase cells counted for each condition). Error bars represent SD of the mean. ns: not significant, ***: *P* ≤ 0.001. (C) Measurement of inter-centromere distances on chromosome 11 in G2 cells after depletion of NSD3 or RAD21. *Bottom*, Representative images of subsequent FISH experiments performed using a specific probe for chromosome 11 centromere (Cen-Chr11). Also shown is a magnified view of one the three loci of Cen-Chr11. (D) Box plot of the distance between paired FISH signals (*n* ≥ 150) for each tested siRNA in three independent experiments, with plots showing the 5 and 95 percentiles. ns: not significant; ****: *P* ≤ 0.0001).

Since NSD3 does not affect regulatory pathways involved in protecting mitotic cohesion, we then explored whether the defective cohesion seen after NSD3 depletion is also detected in interphasic cells before mitotic entry. To this end, we analysed sister chromatid cohesion in G2-synchronised cells by DNA fluorescent *in situ* hybridization (FISH) using a probe specific for the centromeric region of chromosome 11 (Figure 2C). Cohesion was assessed by measuring the distances between paired DNA FISH signals in control cells and in cells depleted for NSD3, with RAD21 depletion used as a positive control for G2 cohesion defects (see Figure S4 for validation of both depletions). The mean distance between paired dots was 0.52 μm in the control cells, and this increased to 0.67 μm upon NSD3 depletion and to 0.68 μm in RAD21-depleted cells (Figure 2D). These results demonstrate that NSD3 depletion leads to defective sister chromatid cohesion in G2 cells, thereby indicating that NSD3’s contribution to cohesion must take place before cells enter mitosis.

We next addressed whether NSD3 could be involved in the loading of cohesin and/or kollerin onto chromatin, a process that takes place at mitotic exit and early in G1 phase. For this purpose, control and NSD3-depleted cells were synchronised by a single thymidine arrest-and-release, followed by mitotic arrest induced by nocodazole (Figure 3A). We harvested the cells at different time points after their release from the nocodazole-mediated arrest, then fractionated them to obtain chromatin fractions. We next analysed these by SDS-PAGE and immunoblotting. Under these experimental conditions, NSD3 inactivation did not impact cell mitotic exit, since both the unloading of the CAP-D2 condensin complex subunit I from chromatin and the dephosphorylation of serine 10 on histone H3 occured with identical kinetics in the control and NSD3-inactivated cells (Figure 3B-C). By contrast, the accumulation of cohesin subunits (RAD21, SA-2, and SMC1A) onto chromatin over time was lower in NSD3-depleted cells than in control cells. Remarkably, a similar decrease was also observed for the kollerin subunit MAU2. These decreased protein signals were not due to reduced amounts of chromatin, as both the chromatin-associated enzyme topoisomerase II and histone H3 were present at comparable levels in the control and NSD3-depleted cells. The effect cannot be attributed to a decrease in total proteins either, because whole-cell extracts had constant protein levels (Figure 3B left panels). This experiment thus showed that NSD3 inactivation leads to reduced recruitment of cohesin and MAU2 onto chromatin during mitotic exit, suggesting that NSD3 contributes to this process.

**Figure 3:**
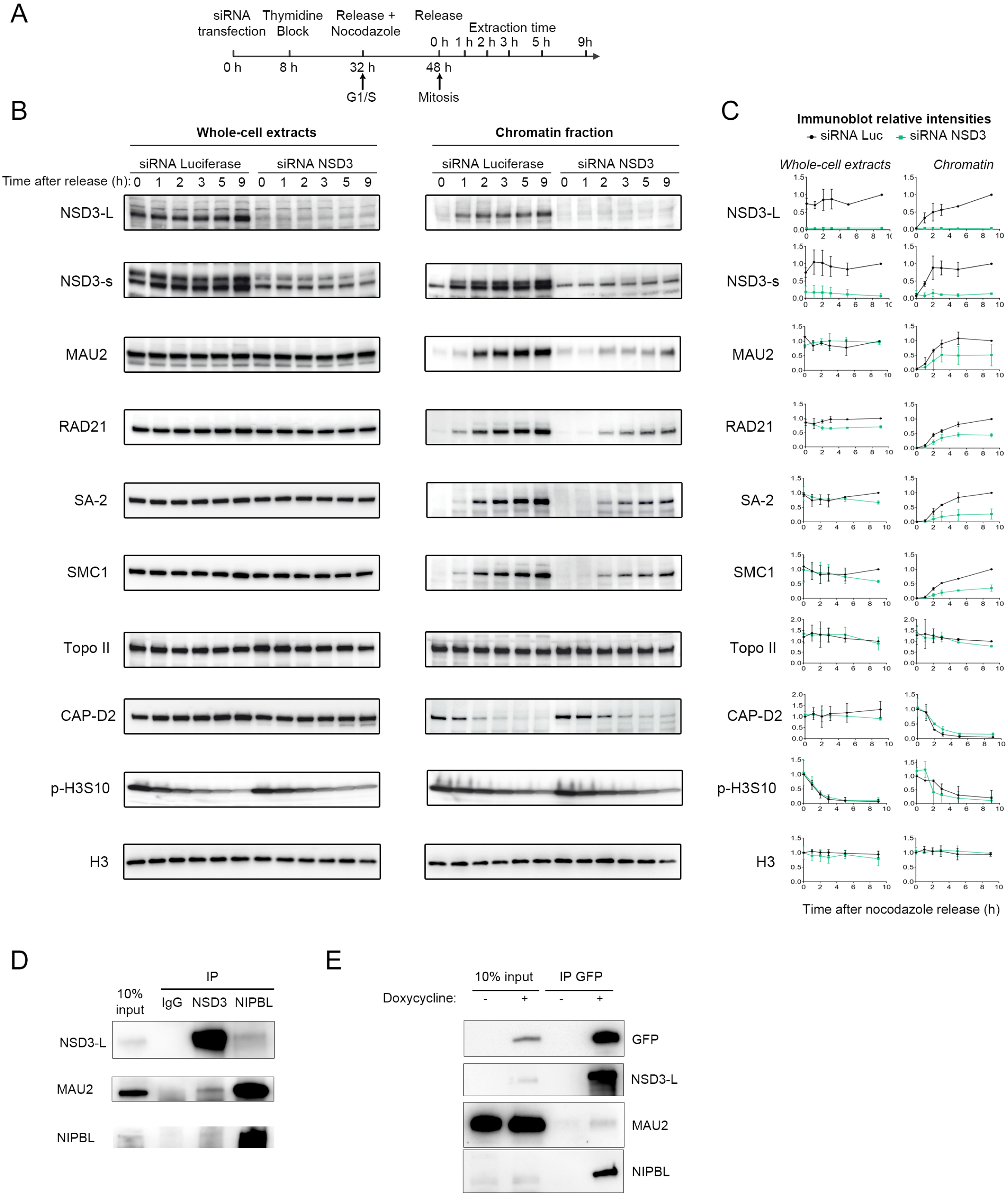
NSD3 interacts with the kollerin complex, favouring cohesin and kollerin loading onto chromatin at mitotic exit. (A) Schematic representation of the synchronization protocol used before harvesting cells for the analysis of MAU2 and cohesin loading after depletion of NSD3. (B) Cohesin loading assay from cells treated with luciferase (Luc) or NSD3 siRNA. Following release from a prometaphase arrest, at the times indicated (in hours), chromatin-bound fractions were prepared and analysed by western blot with the indicated antibodies. Shown are representative blots from one experiment. (C) Signal intensity for each antibody tested in (B) relative to that of time 0 h in the corresponding luciferase-depleted condition (*n* = 2). Error bars represent SD of the mean. (D-E) Co-immunoprecipitation of NSD3 with the kollerin (NIPBL/MAU2) complex. In (D), chromatin extracts from HeLa cells were immunoprecipitated with rabbit IgG (the control), anti-NSD3, or anti-NIPBL antibodies. In (E), chromatin extracts from HeLa EmGFP-NSD3 cells with or without doxycycline treatment were submitted to immunoprecipitation with an anti-GFP antibody. An input corresponding to 10% of the amount of each chromatin extract was loaded, and after immunoprecipitation the eluted fraction was analysed by western blot with antibodies against NSD3, MAU2, and NIPBL. In (E), an additional immunoblot was performed with an anti-GFP antibody to validate the immunoprecipitation of the exogenous EmGFP-NSD3.

Next, we hypothesized that NSD3 interacts physically with the kollerin complex to recruit it to chromatin complexes. Using HeLa cell chromatin extracts, we immunoprecipitated endogenous NSD3, and we observed co-precipitation of both MAU2 and (albeit to a lesser extent) NIPBL (Figure 3D). In a reciprocal experiment using the anti-NIPBL antibody, MAU2 was strongly co-immunoprecipitated as expected, and so was NSD3. To confirm this interaction, we used a cell line that allows for the inducible expression of full-length NSD3 fused to an emerald green fluorescent protein (EmGFP) tag (Figure S5). Using an anti-GFP antibody, we could immunoprecipitated EmGFP-NSD3 together with MAU2 and NIPBL only when EmGFP-NSD3 was induced (Figure 3E). Our results show therefore that NSD3 is able to interact physically with the kollerin complex.

### NSD3 is released from chromatin at mitosis onset then reloaded in early anaphase

As shown in Figure 3B and C, immunoblot analysis of chromatin fractions revealed that both the long and short NSD3 variants are absent from chromatin in mitosis-arrested control cells, and that they are then progressively recruited onto chromatin upon release from mitotic arrest and progression to the G1 phase. This behaviour is similar to that of kollerin and most of cohesin. Our next step was to more precisely describe the dynamics of chromatin loading of NSD3 relative to those of kollerin and cohesin at the single cell level by fluorescence microscopy approaches.

Having checked the suitability of the anti-NSD3 antibody for indirect immunofluorescence experiments (Figure S6), we monitored the localisation of NSD3 at different stages of the cell cycle, using chromosome morphology and H3S10 phosphorylation to identify the different mitotic stages. To assess the presence of NSD3 on chromatin, we performed immunostaining experiments with and without detergent-based pre-extraction of the soluble pool of proteins (Figure S7A-B). As it is the case for many proteins that bind chromatin, we found that NSD3 localizes in the nucleus during interphase, is evicted from chromatin in prophase, then re-loads onto chromatin from anaphase onwards. Finally, to ensure that NSD3 localisation is not altered by fixation, we performed live-cell imaging in a cell line expressing EmGFP-NSD3 and H2B-mCherry, and were able to confirm that EmGFP-NSD3 presents the same localisation profile and dynamics as the ones revealed by immunofluorescence experiment (Supplementary Movies 1-3).

We then analysed the presence of EmGFP-NSD3 on chromatin from metaphase to mitotic exit, comparing its signal to those obtained with anti-RAD21 and anti-MAU2 antibodies, used as cohesin and kollerin markers, respectively. We saw that EmGFP-NSD3 signals were detected on chromatin prior those of MAU2 and RAD21, indicating that NSD3 loads onto chromatin before MAU2 and RAD21 (Figure 4A-B). NSD3 is therefore removed from chromatin after mitosis entry and is reloaded back at early anaphase shortly before kollerin and cohesin are recruited.

**Figure 4:**
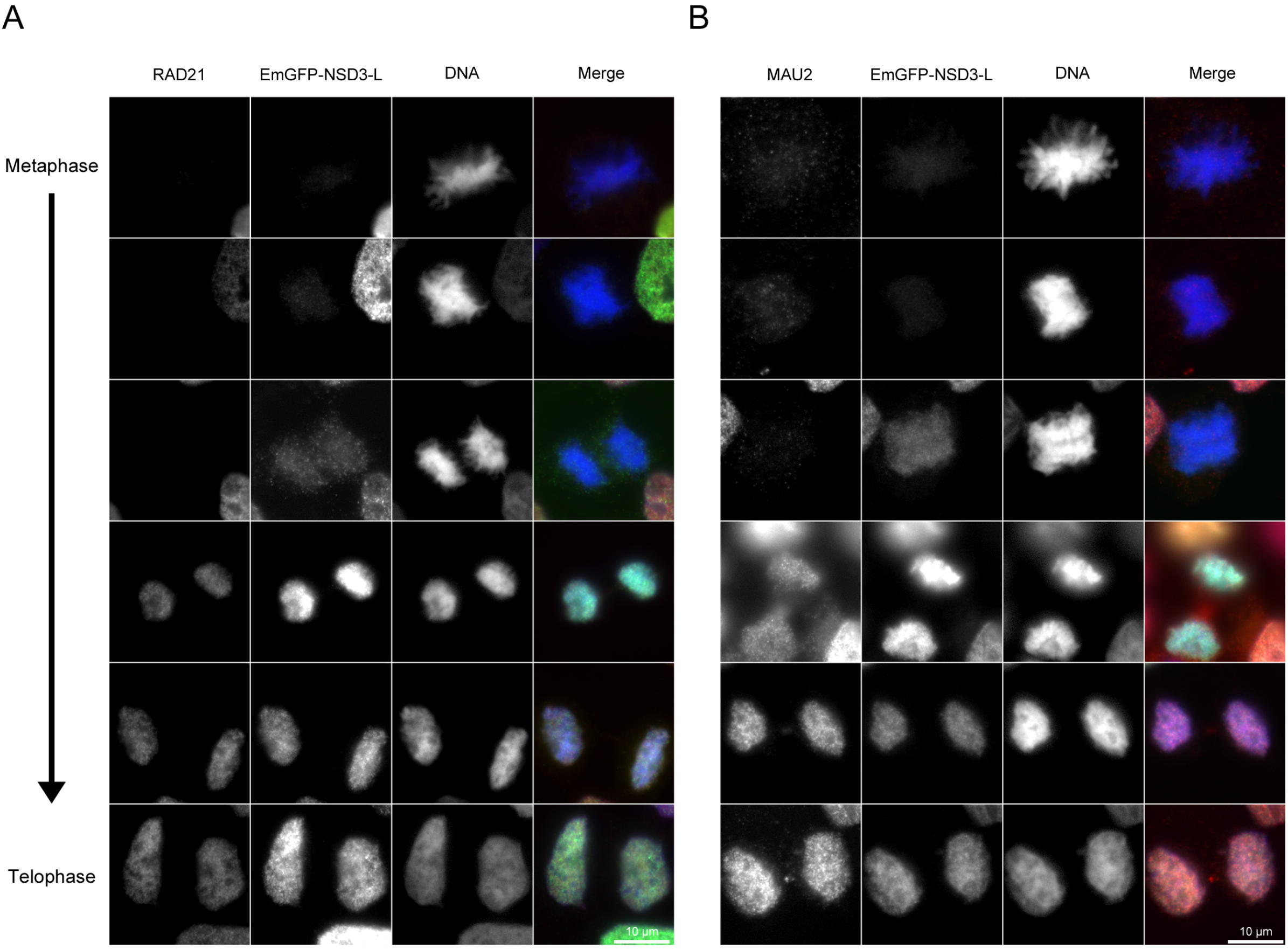
Timing of NSD3, MAU2, and RAD21 recruitment onto chromatin after the metaphase-anaphase transition. Representative images from metaphase to telophase of cells expressing the EmGFP-tagged long isoform of the NSD3 methyltransferase. The cytosol was extracted before immunolabelling with anti-RAD21 (A) or anti-MAU2 (B) antibodies (see Materials and methods).

### Sister chromatid cohesion and cohesin loading are mediated by the long NSD3 isoform and depend on its methyltransferase activity

Our work shows that RNAi-mediated inactivation of NSD3 results in the reduction of cohesin and kollerin loading onto chromatin as well as cohesion defects in both interphase and mitotic cells. As we had used siRNA targeting both the long and short NSD3 isoforms, we decided to explore whether one or both isoforms participate in these processes. To this end, we designed siRNAs directed against the distinct 3’-UTR domain of each isoform (two different siRNAs per isoform). These siRNAs enabled to deplete each form in an efficient and selective manner, as shown by immunoblotting experiments (Figure 5A). We then compared the ability of these isoform-specific siRNAs to induce PSCS in mitotic cells. As shown in Figure 5B, both of the siRNAs targeting the long isoform resulted in defective cohesion, while those targeting the short form did not despite efficient NSD3-s depletion. We next analysed the consequence of isoform-specific depletion on SAC-dependent arrest by using the most efficient siRNA for each isoform, NSD3-s-1 and NSD3-L-2. Depletion of NSD3-s did not have any effect on the levels of phosphorylated H3S10 (Figure 5C). In contrast, when NSD3-L is absent, phosphorylated H3S10 signals accumulate strongly, and is sensitive to a treatment with reversine (Figure 5C). These results show that depletion of only the long form of NSD3 is involved in the accumulation of cells in mitosis as well as in sister chromatid cohesion defects. This strongly suggests that NSD3-L is the one NSD3 isoform involved in cohesion regulation.

**Figure 5:**
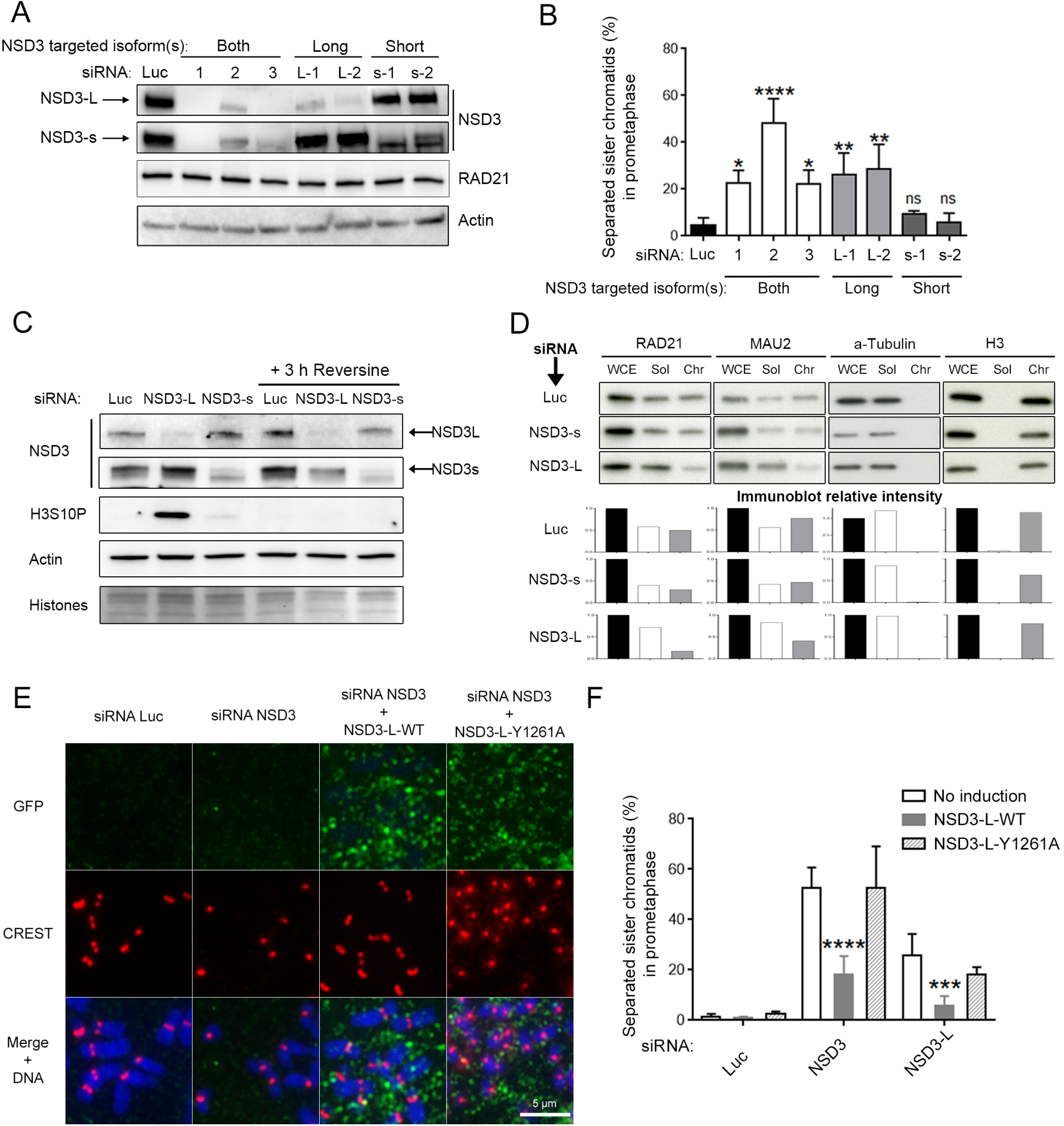
Sister chromatid cohesion and cohesin loading are mediated by the long NSD3 isoform and depend on its methyltransferase activity. (A) Depletion efficiency of the long and short NSD3 isoforms after 72 h transfection with the indicated siRNAs. RAD21 and actin were used as the nuclear and cellular loading markers, respectively. Note that in the luciferase (Luc), NSD3-L-1, and NSD3-L-2 lanes, the NSD3-s band and non-specific band just below it appear to be fused. (B) Percentage of separated sister chromatids in cells depleted for specific NSD3 isoforms during prometaphase (*n* =3, ≥ 150 prometaphase cells counted in each condition). Error bars represent SD of the mean. ns: not significant; *: *P* ≤ 0.05; **: *P* ≤ 0.01: ****: *P* ≤ 0.0001. (C) WB analysis of cyclin B1 and phosphorylated H3S10 mitotic markers after 72 h transfection with siRNA which specifically target the NSD3-s and NSD3-L isoforms. For each condition, half of the cells were treated for 3 h before protein extraction with 1 μM reversine to inhibit the SAC. Note the presence of a non-specific band just below the NSD3-s band. (D) Representative WB of MAU-2 and RAD21 amounts in different cellular fractions obtained from cells depleted for 72 h with siRNA targeting specific NSD3 isoforms. WCE, Sol, and Chr correspond to whole-cell extracts and soluble and chromatin fractions, respectively (see Materials and methods). α-tubulin and histone H3 were used as the cytosolic and chromatin markers, respectively. *Bottom*, For each sample, the signal intensity was quantified. Shown are the ratios calculated relative to that of the whole-cell extracts after the same siRNA treatment. (E-F) Percentage of separated sister chromatids in NSD3-L rescue experiments. To show this, cells were transfected and induced when required with doxycycline for 72 h. They were then immunolabelled with anti-GFP antibodies to improve NSD3 signal detection, and with CREST serum to detect centromeres. (E) Representative images of chromosome spreads from cells transfected with only Luc siRNA, only NSD3 siRNA, or NSD3 siRNA and either EmGFP-NSD3-L (WT) or the catalytic-inactive mutant Y1261A. (F) The percentage of separated sister chromatids in prometaphase cells (*n* =3, 100-300 prometaphase cells counted in each condition). Error bars represent SD of the mean. ***: *P* ≤ 0.001: ****: *P* ≤ 0.0001.

We therefore went on to compare the effect of depleting each NSD3 isoform on the levels of MAU2 and of cohesin bound to chromatin by subcellular fractionation experiments (Figure 5D; also see Figure S8 for isoform-depletion efficiency). Fractionation efficiency was verified with α-tubulin, a cytosolic protein present in the soluble fraction but not in the chromatin-associated fraction, and with histone H3, a core component of chromatin which is not detected in the soluble fraction. As expected, RAD21 and MAU2 are present in both soluble and chromatin fractions in the control cell extracts. However, depletion of NSD3-L induced a decrease of MAU2 and RAD21 amounts in the chromatin fraction while, at the same time, these are conversely increased in the soluble fractions (Figure 5D). This supports the hypothesis that the NSD3 long isoform (but not the short NSD3-s) is the one that contributes to cohesin loading.

In order to further strengthen this view and exclude any off-target effect that siRNAs may have, we tested whether expression of exogenous NSD3-L could rescue defective sister chromatid cohesion induced by depletion of its endogenous counterpart. For this purpose, we performed rescue experiments using the cell line expressing EmGFP-NSD3-L with the siRNA NSD3-2, which targets both isoforms, as well as with NSD3-L-2, which is specific for the long variant only. We used western blots to verify proper expression of exogenous EmGFP-NSD3-L and efficient depletion of endogenous NSD3 isoforms (Figure S9). Then, we determined the percentage of prometaphase cells exhibiting PSCS for each condition. To take into account only EmGFP-NSD3-L-positive cells in rescue experiments and to detect sister chromatid separation concomitantly, we deposited swollen cells onto a glass slide with a cytocentrifuge from the same populations used for western blots, and then labelled NSD3 and centromere by immunofluorescence with an anti-GFP antibody and a CREST serum, respectively. As expected since NSD3 is not associated with chromatin during prometaphase, EmGFP-NSD3-L was diffused around the chromosome (Figure 5E). In the absence of doxycycline, we confirmed that depletion of both NSD3 isoforms or of just the long form results in PSCS (Figure 5F). By contrast, in cells where wild-type EmGFP-NSD3-L was expressed, the proportion of mitotic cells displaying PSCS was significantly reduced (Figure 5F). This indicates that ectopic correction of NSD3-L protein level rescues defective mitotic cohesion to a large extent, thereby establishing NSD3-L as the isoform required for proper sister chromatid cohesion.

Finally, we addressed whether NSD3 methyltransferase activity is required for sister chromatid cohesion. To this end, we generated another cell line expressing a catalytically inactive NSD3-L mutant by substituting tyrosine 1261 to an alanine inside the catalytic pocket of the SET domain^58^. When the same rescue experiment as the one described above was performed with these cells, expression of the EmGFP-NSD3-Y1261A mutant did not attenuate the cohesion defects induced by endogenous NSD3 depletion (Figure 5F). Therefore, the methyltransferase activity of the long NSD3 isoform is required to ensure regulation of sister chromatid cohesion.

## Discussion

Sister chromatid cohesion is essential for proper chromosome segregation. In this work, we show that depletion of the methyltransferase NSD3 leads to a defect in this process and that this is characterized by a mitotic delay dependent on the spindle assembly checkpoint. Because the key WAPL regulator is unable to rescue PSCS induced by depletion of NSD3, we ruled out the possibility that NSD3’s role in cohesion could be related to processes involving cohesin dissociation or cohesion maintenance during mitosis. Because of its involvement in kollerin recruitment and in cohesin loading onto chromatin, we propose that defective kollerin recruitment during mitotic exit is the cause of the PSCS observed in mitotic cells upon NSD3 inactivation. In agreement with this interpretation, the defects observed in mitotic sister cohesion are incomplete, and depending on the siRNA used involve 20-40% of the cells, being consistent with the cohesion defect levels usually observed in vertebrate cells with depletion of kollerin or of other proteins known to be involved in cohesin loading during mitotic exit^5,59,60^. However, cohesin loading occurs not only at mitotic exit, but also in S-phase^26^. In that case, cohesin loading is dependent on Mcm2-7 and involves kollerin, whose alteration leads to cohesion defects before mitotic entry. Thus, the mitotic cohesion defects observed when NSD3 is absent may also be attributed to its possible contribution to kollerin loading during the pathway specific to the S-phase. Moreover, because NSD3 depletion is likely to have pleiotropic effects due to its role in chromatin modification, we cannot definitively rule out the hypothesis that NSD3 is also involved in other interphasic cohesion-related processes.

Looking for a NSD3’s contribution in cohesion during interphase, we show that NSD3 interacts with kollerin on chromatin. We also show that NSD3 is required for the recruitment of MAU2 to chromatin during mitotic exit. As a consequence, NSD3 also contributes to the loading of cohesin onto chromatin. In full agreement with this function, we showed that NSD3 recruitment to chromatin occurs during early anaphase, slightly before those of MAU2 and RAD21^5^. The physical interaction we uncovered between NSD3 and kollerin is therefore unlikely to occur prior to their targeting to chromatin. Instead, NSD3 may promote the binding of kollerin to chromatin during mitotic exit. Consistent with this possibility, the MAU2 homologue in budding yeast, Scc4, is necessary for loading Scc2 onto chromatin *in vivo*, but it has no affinity for DNA *per se*, implying that a protein receptor is required^10–12^ It has also been reported that the transcription regulator BRD4 co-immunoprecipitates with NSD3 and with the kollerin complex through its extra-terminal domain, stabilizing NIPBL on chromatin in vertebrate cells^46^. In RN2 human acute myeloid leukaemia cells, BRD4 recruits NSD3 on chromatin at active promoters and enhancers across the genome^61^, and these domains are known to be enriched with cohesin^62–64^ In *Drosophila melanogaster*, the BRD4 homologue Fs(1)h promotes the association of Nipped-B and RAD21 with enhancers and promoter regions close to the replication origin, and genetic interactions exist between genes which code for these three proteins^65^. It is thus tempting to propose that BRD4 and NSD3 act together to load kollerin onto chromatin. However, we showed that NSD3 depletion decrease RAD21 level on chromatin and induced PSCS that are not rescued by WAPL depletion. By contrast, the absence of BRD4 do not decrease the level of RAD21 on chromatin in mouse embryonic stem cells and induces differentiation defects in neural crest progenitors that are rescued by WAPL depletion^66^. Moreover, the link between NSD3 and BRD4 depends on the short NSD3 isoform at the functional level^61^, whereas we have demonstrated that the long NSD3 isoform is responsible for the regulation of sister chromatid cohesion. Thus, the putative interplay between BRD4 and NSD3 in chromatin organisation remains to be clarified. Future studies will also be necessary to precise the molecular mechanism of NSD3-dependent kollerin recruitment, and notably to confirm its specificity, to elucidate whether it occurs through direct interaction with the kollerin complex or by providing a chromatin context which is suitable for its binding. Interestingly, it has already been reported that NSD3 can play such adaptor role, bridging the BET domain of BRD4 and the CHD8 chromatin remodeling factor^54^ In *Schizosaccharomyces pombe*, the chromatin remodeling complex RSC (Remodels the Structure of Chromatin) acts as a chromatin receptor by physically interacting with the Scc2/Scc4 complex. It also promotes nucleosome eviction in order to render the DNA naked, thus facilitating cohesin loading^67–70^. Moreover, other chromatin remodelers have been described as contributing to cohesin loading^64,65,70–73^. Looking for a link between NSD3 and chromatin remodeling is therefore a promising way to explore the contribution of NSD3 to kollerin chromatin-association.

By targeting specific isoforms, we were able to show that the NSD3’s contribution to sister chromatid cohesion and to chromatin-association of kollerin and cohesin actually only involves the full-length form which contains the methyltransferase activity. Moreover, we also showed that unlike its wild-type counterpart, a catalytically inactive version of NSD3-L was not able to rescue cohesion defects. These results imply that NSD3-L regulates sister cohesion by methylating one or more substrates that have not yet been identified. Interestingly, the methyltransferase Suv4-20h2 was previously shown to specifically regulate cohesin loading on pericentromeric heterochromatin, known to have low transcriptional activity^59^. This suggests that complementary cohesin-loading regulating pathways do co-exist according to the targeted chromatin domains. Because kollerin seems to require partners for its chromatin interactions^10–12^, one can speculate that the catalytic activity of NSD3-L might act to recruit kollerin at specific chromatin sites, which is consistent with the partial defects observed in kollerin loading and in mitotic sister cohesion. As for right now, the histone H3K36 is the only described substrate of NSD3. Thus, H3K36me2 stands as an attractive candidate to further investigate NSD3’s contribution to the recruitment of cohesin loaders and cohesin onto chromatin. In agreement with this possibility, H3K36me2 is found in active gene promoters, where NIPBL is also enriched in human cells^74–76^. Such a methylated mark which is stable throughout cell division could be essential for the epigenetic inheritance of some cohesin loading sites.

## Materials and methods

### Antibodies

Primary antibodies used in this work with dilutions for western blotting (WB) and immunofluorescence (IF) are indicated here:

| Antibody | Host | Source | Identifier | WB dilution | IF dilution |
| --- | --- | --- | --- | --- | --- |
| NSD3 | Rabbit | ProteinTech | 11345-1-AP | 1/1000 | 1/1000 |
| RAD21 | Mouse | Millipore | 05-908 | 1/5000 | 1/1000 |
| MAU2 | Rabbit | Abcam | Ab183033 | 1/5000 | 1/1000 |
| NIPBL | Rabbit | Bethyl Laboratories | A301779A |  |  |
| SA-2 | Mouse | Santa Cruz Biotechnology | sc-81852 | 1/500 |  |
| SMC3 | Rabbit | Ref : <sup>77</sup> |  | 1/1000 |  |
| WAPL | Rabbit | Ref : <sup>16</sup> |  | 1/1000 |  |
| ESCO2 | Rabbit | Ref : <sup>78</sup> |  | 1/1000 |  |
| Sororin | Mouse | Abcam | Ab168217 | 1/1000 |  |
| hCAP-D2 | Rabbit | Ref: <sup>79</sup> |  | 1/1000 |  |
| SGO1 | Rabbit | Ref: <sup>80</sup> |  |  | 1/1000 |
| Topoisomerase IIα (Ki-S1) | Mouse | | MAB4197 | 1/2000 | |
| Phospho-Histone H3(S10), clone 3H10 | Mouse | Millipore | 05-806 | 1/10000 | 1/20000 |
| Cyclin B1 | Mouse | Santa Cruz Biotechnology | Sc-245 | 1/1000 |  |
| Histone H3K36me2 | Rabbit | Active Motif | 39056 | 1/5000 |  |
| Histone H3 | Rabbit | Sigma-Aldrich | H0164 | 1/10000 |  |
| Actin | Rabbit | Sigma-Aldrich | A5060 | 1/10000 |  |
| α-Tubulin | Mouse | Sigma-Aldrich | T5168 | 1/5000 | |
| GFP | Mouse | Roche | 11814460001 | 1/500 | 1/500 |
| CREST serum | Human | Dr Bahon<br>Riedinger;<br>CHRU<br>RENNES |  |  | 1/4000 |

Horseradish peroxidase coupled secondary antibodies (Jackson Immunoresearch) were used at 1/5000 and 1/25000 dilution for WB detection of mouse and rabbit antibodies, respectively. Alexa Fluor-coupled secondary antibodies from Invitrogen were used at 1/1000 dilution for IF detection.

### Plasmid construction and cell line generation

A fragment containing an inducible TRE-tight promoter and a LAP tag (a FLAG-tag followed by an EmGFP-tag) was inserted at the XhoI site of the plasmid pBSKDB-CAG-rtTA2sM2-IRES-tSkid-IRES-Neo (Addgene #62346), thereby generating the vector pGEH_ind_LAP-C. In this vector, the coding sequences of NSD3-s, NSD3-L, and NSD3-L-Y1261A were each cloned in-frame in the C-terminal of the LAP tag after PCR amplification from the plasmids pMSCV_MigR1_NSD3short and pMSCV_MigR1_NSD3long (both kindly provided by Prof. C.R. Vakoc)^54^. For the construction of NSD3-L-Y1261A, the canonical codon sequence “tat” generating Y1261 was replaced by “gct” on the PCR fragments used for the cloning reaction, leading to the generation of A1261. All PCR reactions were done using Q5 high-fidelity polymerases (M0493S, NEB), and all constructions and intermediates were generated using a NEBuilder HiFi DNA assembly cloning kit (E5520S, NEB) according to the manufacturer’s instructions. Plasmids and their maps can be provided upon request.

For generation of HeLa EmGFP-NSD3-L, and EmGFP-NSD3-L-Y1261A cell lines, 5 μg of the corresponding plasmids supplemented with 10 μl of P3000 in 125 μl Opti-MEM (31985062, Thermo Fisher Scientific), was mixed with 7.5 μl Lipofectamine 3000 reagent in 125 μl Opti-MEM. After 5 min incubation at room temperature, the plasmid-containing mix was transfected in HeLa Kyoto cells seeded the day before in 2.25 ml of complete medium in 6-well plates. After 24 h, the cells were trypsinized and either 1/10 or 1/50 of the cells were seeded in culture dishes (150 mm) with 25 ml of complete medium containing 1 mg/ml of geneticin selective antibiotic (11811023, Thermo Fisher). Selection lasted for 2-3 weeks, with replacement of the antibiotic-supplemented medium every 2-3 days. The cells were then induced for EmGFP-NSD3 expression with 2 μg/ml of doxycycline (D9891-1G, Merck) for 48 h, and GFP-positive cells were FACS-sorted at the Biosit CytomeTRI platform. They were cultured for another 2 weeks under selection pressure but without induction, and the cell lines were then subcloned by limiting dilution in 96-well plates. After 2 weeks of selection, EmGFP-NSD3 expression was evaluated in about 30 clones for each NSD3 variant. Despite this process, no homogeneous clonal cell lines could be obtained. For the EmGFP-NSD3-s construct, the clones obtained always expressed the fused protein constitutively. In contrast (and as expected), expression of EmGFP-NSD3-L and EmGFP-NSD3-L-Y1261A in the selected cell lines was inducible. A HeLa H2B-mCherry EmGFP-NSD3-L cell line was generated by transfecting EmGFP-NSD3-L cells with a pH2B_mCherry_IRES_puro2 plasmid (Addgene #21045) as described above, with a 15-day selection and 0.5 μg/ml puromycin.

### Cell cultures and treatment

All experiments discussed in this paper were performed with HeLa Kyoto cells or cell lines constructed from those cells. All cell lines were cultured at 37 °C in an incubator supplied with 5% CO_2_ in Dulbecco’s Modified Eagle Medium (DMEM) with glutamine analogue GlutaMAX (31966047, Thermo Fisher), supplemented with 10% foetal bovine serum (FBS) and a cocktail of penicillin/streptomycin antibiotics (15140-122, Thermo Fisher) at 100 U/ml and 100 μg/ml final concentrations, respectively. Stable cell lines were maintained in culture with 1 mg/ml geneticin. Doxycycline was used at 1 μg/ml in medium to induce expression of the exogenous tagged proteins, and was replaced with fresh doxycycline-containing medium after 48 h for experiments requiring 72 h induction. For SAC experiments, cells were incubated with 1 μM reversine (S7588, Selleck Chemicals) for 3 h or with 100 ng/ml nocodazole (M1404, Merck) for 6 h. To synchronize cells, thymidine (T1895-1G, Merck) was added to the medium 8 h post-transfection at a final concentration of 2 mM, and the cells were cultured for 24 h. To arrest the cells in prometaphase, this was followed by two successive 3-min PBS washes then incubation with complete medium supplemented with 100 ng/ml nocodazole for another 16 h. Cells were harvested by mitotic shake-off, washed as described for the previous release, and seeded in fresh complete medium on new plates. To enrich the cells in G2, they were subjected to a double thymidine block (16 h each time), followed by a release of 6 h. Before use in synchronisation experiments, all media were pre-warmed to 37 °C.

### siRNA transfection

Unless otherwise stated, HeLa cells were transfected with 20 nM siRNA for 72 h with Qiagen HiPerFect reagent according to the manufacturer’s recommendations. As a general guideline, for a 6-well transfection, 10 μl of reagent and 2.5 μl of 20 μM stock siRNA were mixed in 87.5 μl of Opti-MEM. This was added to 300,000 HeLa cells suspended in 2.4 ml of complete medium. Media was replaced after 48 h of transfection, and cells were trypsinized and diluted if necessary. For WAPL co-depletion, cells were transfected with Luc or the indicated siRNA, then 24 h later a second transfection was performed with Luc or WAPL siRNA, and cell culture was prolonged for 48 h. The siRNA used are described below, and were purchased from Qiagen or Dharmacon. A listing of those used for methyltransferase screening can be provided upon request.

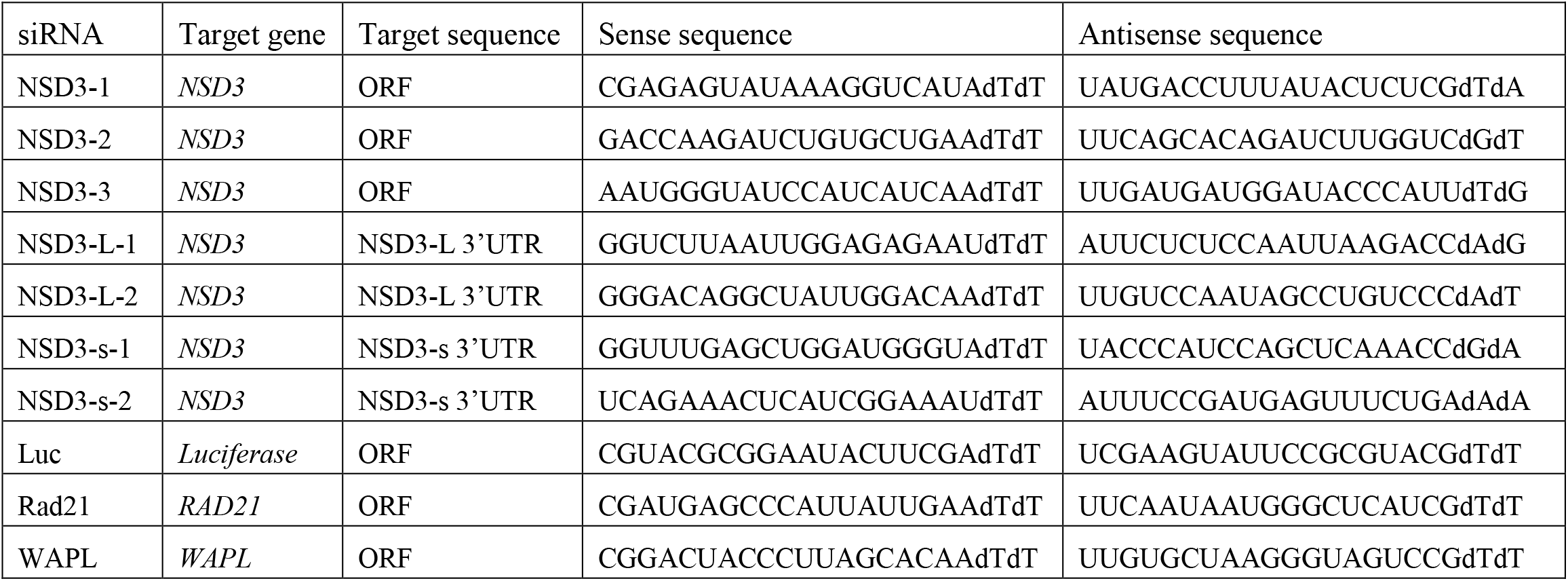

### Chromosome spreads

Cells harvested from a 6-well plate were subjected to hypotonic shock in 4.5 ml of 75 mM KCl for 15 min at room temperature, then 500 μl of Carnoy’s fixative solution (at a ratio of 3:1 methanol to acetic acid) was added. Cells were centrifuged at 250*g* and the pellet was resuspended in 5 ml of the fixative. This operation was repeated three more times and the pellet was kept overnight at −20 °C in 300 μl of Carnoy’s fixative. We then dropped 30 μl of cells onto a dry slide, let them dry for 2 h at room temperature, and incubated the slides for 5 min in fresh 5% GIEMSA solution (1.09203, Merck) diluted in 100 ml Gurr buffer (10582-013, Thermo Fisher). After 3-5 washes in distilled water, Giemsa-stained cells were mounted with Entellan. For immunofluorescence in chromosome spreads, 50,000 cells swollen after hypotonic shock in 200 μl of KCl 75 mM were cytospun on a slide for 5 min at 900 rpm (Cytospin 4, Thermo Fisher). These cells were then fixed in 3% paraformaldehyde/PBS for further immunolabelling.

### Immunofluorescence

When stated, soluble contents of cells were pre-extracted before fixation by incubation for 1 min in 0.1% Triton (T8787-50ML, Merck) diluted in PBS 1X (10010-015, Thermo Fisher). Cells were fixed 10 min in 4% paraformaldehyde (15710, Electron Microscopy Science) diluted in PBS 1X (pH 7.2-7.4). Slides or coverslips were then washed three times for 5 min in PBS 1X, permeabilized with 0.1% Triton X-100 for 10 min, washed three times in PBS 1X, then blocked by incubation with 5% FCS in PBS 1X for 1 h at room temperature. This last solution was used to dilute primary and secondary antibodies. Slides or coverslips were then incubated overnight at 4 °C with primary antibodies, washed three times with PBS 1X, and further incubated for 1 h at room temperature with fluorochrome-conjugated secondary antibodies. The DNA was stained with DAPI (100 ng/ml in PBS), and slides were mounted in ProLong Gold medium (P36982, Thermo Fisher).

### Microscopy and image analysis

Images were acquired with a Zeiss Axio Imager M2 epifluorescence microscope equipped with Zeiss Plan-Apochromat 40x/1.3 and 63x/1.40 oil objectives, a CoolSNAP HQ^2^ CCD camera (Photometrics), and Zeiss AxioVision software (version 4.2). Signals were quantified using the Fiji image processing package^81^.

### Fluorescent in situ hybridization (FISH)

DNA FISH was performed as previously described^82^, except that cells were fixed with Carnoy’s fixative (as described above), then spread on glass sides before being processed. Analysis was only done on pairs for which the dots could be clearly resolved. The probe used for FISH was coupled with an Alexa 488 fluorophore and targets the alpha-satellite sequence AgGgTtTcAgAgCtGcTc specific to the centromeric region of chromosome 11. In that probe sequence, uppercase letters correspond to DNA, while lowercase letters indicate locked nucleic acids, which are modified RNA nucleotides in which the ribose moiety is modified, with an extra bridge connecting the 2’ oxygen and 4’ carbon.

### Live cell imaging

HeLa H2B-mCherry EmGFP-NSD3-L cells were seeded on a Lab-Tek chamber slide (155409, Thermo Fisher) in 500 μl DMEM complete medium supplemented with 2 μg/ml doxycycline, then incubated overnight as described above. Afterwards they were supplemented with 10 μM of the CDK1 inhibitor RO-3306 (S7747, Selleck Chemicals) and incubated for 6 h. Cells were washed three times with DMEM preheated to 37 °C, then the medium was replaced with 500 μl of Leibovitz’s L-15 medium (11415064, Thermo Fisher), also preheated to the same temperature and supplemented with 20% FBS, penicillin/streptomycin antibiotics (as above), and 2 mM L-glutamine (25030081, Thermo Fisher). Cells were imaged at 37 °C every 3 min for 10 h, with 15 stacks spaced by 1 μm at each acquisition time. They were irradiated with lasers to excite H2B-mCherry (561 nm) and EmGFP-NSD3-L (488 nm). Images were acquired using a spinning disk system consisting of a Leica DMi8 microscope equipped with a 63X/1.4 oil objective, a CSU-X1 spinning-disk unit (Yokogawa), and an Evolve EMCCD camera (Photometrics). The microscope was controlled using its dedicated software and the Inscoper imaging suite.

### Cell extracts and western blotting

For whole-cell extracts, proteins were extracted by directly resuspending cell pellets in Laemmli buffer (60 mM Tris-HCl pH 6.8, 10% glycerol, 2% SDS, 0.05% bromophenol blue, and 5% β-mercaptoethanol). For fractionation, cells were collected by trypsinization and washed once with ice-cold PBS. The final cell pellet was resuspended in extraction buffer (20 mM Tris pH 7.5, 100 mM sodium chloride, 5 mM magnesium chloride, 0.2% NP-40, 10% glycerol, and 0.5 mM dithiothreitol) supplemented with EDTA-free protease inhibitor tablets (5892953001, Merck) and a homemade phosphatase inhibitor cocktail (final concentrations of 5 mM sodium fluoride, 10 mM β-glycerophosphate, 1 mM sodium pyrophosphate, and 0.2 mM sodium orthovanadate). Cells were lysed on ice by ten passages through a 27-gauge needle. The resulting lysates were incubated for 10 min on ice, an aliquot was collected as a whole-cell extract, and the remaining lysates were centrifuged at 12000*g* for 5 min at 4 °C in order to collect the soluble protein extracts. The chromatin-containing pellet was washed four times with the same extraction buffer at 12,000 g for 5 min at 4 °C and then resuspended in Laemmli buffer.

For immunoblotting, lysed cells were heated for 5 min at 95 °C. Samples were then subjected to SDS-PAGE electrophoresis in a 4-20% polyacrylamide gradient gel (4561094, Biorad) and a Biorad Trans-Blot Turbo transfer system was used to transfer them to PVDF membranes (1704156, Biorad). Following saturation for 1 h in PBS 1X containing 5% milk and 0.1% Tween 20 (655204-100ML, Merck), the membranes were incubated overnight at 4 °C with primary antibodies, then for 1 h at room temperature with horseradish peroxidase (HRP) conjugated secondary antibodies, according to standard procedures. For NSD3 western blot analysis, saturation and incubation were performed similarly, except that the saturation and incubation buffers were supplemented with 10% (instead of 5%) milk and with 150 mM NaCl. Membranes were treated with the Immobilon ECL Ultra Western HRP substrate (WBKLS0500, Merck), and signals were detected using an Amersham Imager 680 (GE Healthcare) or using Amersham Hyperfilm ECL film (GE28-9068-35, Merck). Quantification of band signal intensity was done with Fiji’s analyse gels tool.

### Immunoprecipitation

Cells were resuspended at 4.10^7^ cells/ml of buffer A (10 mM HEPES pH 7.9, 10 mM potassium chloride, 1.5 mM magnesium chloride, 0.34 M sucrose, 10% glycerol, and 0.5 mM dithiothreitol), to which the same volume of buffer A supplemented with 0.2% NP-40 was then added. After 10 min incubation on ice, the extracts were centrifugated at 1,300*g* for 5 min at 4 °C. The pellets were washed for 1 min in buffer A to a concentration equivalent to 20,000 cells/ml. After a similar centrifugation, they were resuspended to 40,000 cells/ml in IP buffer (10 mM HEPES pH 7.9, 10 mM potassium chloride, 140 mM sodium chloride, 1.5 mM magnesium chloride, 10% glycerol, and 0.5 mM dithiothreitol). The lysate was supplemented with calcium chloride to a final concentration of 3 mM and chromatin was solubilised for 2 h at 4 °C by digestion with Turbonuclease (T4330, Merck) to a final concentration of 250U/ml. The reaction was then arrested by addition of a 25X stop buffer (125 mM EGTA and 10 mM EDTA) to a final concentration of 1X. Solubilised chromatin was collected by centrifugation at 13,000*g* for 20 min at 4 °C. For immunoprecipitation against endogenous proteins, 250 μl of chromatin extracts were incubated overnight with 5 μg of IgG, anti-NSD3, or anti-NIPBL rabbit antibodies followed by 1 h incubation at 4 °C on a rotating wheel with 40 μl of a 1:1 mix of Dynabead protein A (10001D, Life Technologies) and G (10003D, LifeTechnologies) magnetic beads. Beads were then washed four times in 500 μl of IP buffer without protease inhibitors, and eluted in 80 μl of Laemmli buffer. For immunoprecipitation of the induced EmGFP-NSD3-L proteins, the protocol was the same except that chromatin solubilisation with Turbonuclease was performed for 1 h only, and was not arrested with the stop buffer. Instead, 25 μl of ChromoTek GFP-Trap M-270 magnetic particles (gtd-20, Proteintech) were immediately added to 250 μl of the extracts. This was incubated for 1 h on a rotating wheel at 4 °C, and before elution was washed four times in 500 μl ice-cooled wash buffer (10 mM Tris/HCl pH 7.5, 150 mM NaCl, 0.05% NP-40, and 0.5 mM EDTA). All buffers except the wash and elution buffers were supplemented with protease inhibitors.

### Statistics

All statistical analysis was performed using GraphPad Prism software (v6.05). For the comparison of the means of the proportions showing PSCS or the mitotic index between NSD3-depleted cell lines, we performed one-way analysis of variance (ANOVA), or in the case of Figure 2B, two-way ANOVA. These were followed by Dunnett *post-hoc* analysis, assuming the normal law for the repartition of the means. For Figures S3A and S3B, we performed a chisquare analysis of the experiment with Yate’s correction. Comparisons of fluorescence intensity from representative experiments were analysed using a nonparametric Kruskal-Wallis test followed by Dunn’s multiple comparison correction. For distance comparison in the DNA FISH experiments, one-way ANOVA was followed by Bonferroni’s multiple comparisons test. For all tests, an alpha risk of 0.05 was used.

## Author contributions

G.E.-H and L.M.-J designed and performed the experiments. G.B contributed to plasmid constructions. G.E-H, L.M-J, and E.W analysed data. G.E-H and E.W wrote the manuscript. G.E-H, E.W and C.J. designed and supervised the project. F.S, R.G and C.J provided funding and revised the manuscript.

## Acknowledgments

We would like to give special thanks to Dr. Rozenn Gallais for her contributions to this study. For providing reagents, we thank Prof. Christopher Vakoc (Cold Spring Harbor Laboratory, USA), Dr. Christophe Escudé (National Museum of Natural History, France), Prof. Jan-Michael Peters (Research Institute of Molecular Pathology, Austria), Prof. Yoshinori Watanabe (University of Tokyo, Japan), and Dr. Isabelle Bahon-Riedinger (Rennes University Hospital, France). We also thank Laurent Deleurme of the CytomeTRI flow-cytometry platform at BIOSIT (SFR UMS CNRS 3480 – INSERM 018) for the FACS cell sorting of the EmGFP-NSD3 HeLa cell lines. We thank Juliana Berland for suggestions on the manuscript.

## Affiliation and fundings

G.E.-H, L.M.-J, G.B, E.W and R.G. are affiliated with the CNRS, and C.J. to INSERM. E.W was supported by grants from Rennes Métropole, the Institut National du Cancer [PLBIO2012], and the European ERA-NET via the E-RARE 2 rare disease program [TARGET-CdLS]. This work was funded by the French National Research Agency [ANR project EpiCentr], the Cancéropôle Grand Ouest, the Fondation pour la Recherche Médicale, and the Ligue contre le cancer, comité du Grand-Ouest. E.W and C.J. were also supported by the Region Bretagne.

## References

1. Nasmyth, K., and Haering, C.H. (2009). Cohesin: its roles and mechanisms. Annu Rev Genet 43, 525–558. 10.1146/annurev-genet-102108-134233 [doi].

2. Makrantoni, V., and Marston, A.L. (2018). Cohesin and chromosome segregation. Curr Biol 28, R688–R693. 10.1016/j.cub.2018.05.019.

3. Nishiyama, T. (2019). Cohesion and cohesin-dependent chromatin organization. Curr Opin Cell Biol 58, 8–14. 10.1016/j.ceb.2018.11.006.

4. Ciosk, R., Shirayama, M., Shevchenko, A., Tanaka, T., Toth, A., Shevchenko, A., and Nasmyth, K. (2000). Cohesin’s binding to chromosomes depends on a separate complex consisting of Scc2 and Scc4 proteins. Mol Cell 5, 243–254. 10.1016/s1097-2765(00)80420-7.

5. Watrin, E., Schleiffer, A., Tanaka, K., Eisenhaber, F., Nasmyth, K., and Peters, J.M. (2006). Human Scc4 is required for cohesin binding to chromatin, sister-chromatid cohesion, and mitotic progression. Curr Biol 16, 863–874.

6. Litwin, I., and Wysocki, R. (2018). New insights into cohesin loading. Curr Genet 64, 53–61. 10.1007/s00294-017-0723-6.

7. Gillespie, P.J., and Hirano, T. (2004). Scc2 couples replication licensing to sister chromatid cohesion in Xenopus egg extracts. Curr Biol 14, 1598–1603.

8. Takahashi, T.S., Yiu, P., Chou, M.F., Gygi, S., and Walter, J.C. (2004). Recruitment of Xenopus Scc2 and cohesin to chromatin requires the pre-replication complex. Nat Cell Biol 6, 991–996. 10.1038/ncb1177.

9. Tonkin, E.T., Wang, T.J., Lisgo, S., Bamshad, M.J., and Strachan, T. (2004). NIPBL, encoding a homolog of fungal Scc2-type sister chromatid cohesion proteins and fly Nipped-B, is mutated in Cornelia de Lange syndrome. Nat Genet 36, 636–641. 10.1038/ng1363.

10. Chao, W.C., Murayama, Y., Muñoz, S., Costa, A., Uhlmann, F., and Singleton, M.R. (2015). Structural Studies Reveal the Functional Modularity of the Scc2-Scc4 Cohesin Loader. Cell Rep 12, 719–725. 10.1016/j.celrep.2015.06.071.

11. Hinshaw, S.M., Makrantoni, V., Kerr, A., Marston, A.L., and Harrison, S.C. (2015). Structural evidence for Scc4-dependent localization of cohesin loading. Elife 4, e06057. 10.7554/eLife.06057.

12. Murayama, Y., and Uhlmann, F. (2014). Biochemical reconstitution of topological DNA binding by the cohesin ring. Nature 505, 367–371. 10.1038/nature12867.

13. Parenti, I., Diab, F., Gil, S.R., Mulugeta, E., Casa, V., Berutti, R., Brouwer, R.W.W., Dupé, V., Eckhold, J., Graf, E., et al. (2020). MAU2 and NIPBL Variants Impair the Heterodimerization of the Cohesin Loader Subunits and Cause Cornelia de Lange Syndrome. Cell Rep 31, 107647. 10.1016/j.celrep.2020.107647.

14. Chan, K.L., Roig, M.B., Hu, B., Beckouët, F., Metson, J., and Nasmyth, K. (2012). Cohesin’s DNA exit gate is distinct from its entrance gate and is regulated by acetylation. Cell 150, 961–974. 10.1016/j.cell.2012.07.028.

15. Gerlich, D., Koch, B., Dupeux, F., Peters, J.M., and Ellenberg, J. (2006). Live-cell imaging reveals a stable cohesin-chromatin interaction after but not before DNA replication. Curr Biol 16, 1571–1578. 10.1016/j.cub.2006.06.068.

16. Kueng, S., Hegemann, B., Peters, B.H., Lipp, J.J., Schleiffer, A., Mechtler, K., and Peters, J.M. (2006). Wapl controls the dynamic association of cohesin with chromatin. Cell 127, 955–967.

17. Lopez-Serra, L., Lengronne, A., Borges, V., Kelly, G., and Uhlmann, F. (2013). Budding yeast Wapl controls sister chromatid cohesion maintenance and chromosome condensation. Curr Biol 23, 64–69. 10.1016/j.cub.2012.11.030.

18. Ouyang, Z., Zheng, G., Song, J., Borek, D.M., Otwinowski, Z., Brautigam, C.A., Tomchick, D.R., Rankin, S., and Yu, H. (2013). Structure of the human cohesin inhibitor Wapl. Proc Natl Acad Sci U S A 110, 11355–11360. 10.1073/pnas.1304594110.

19. Ouyang, Z., Zheng, G., Tomchick, D.R., Luo, X., and Yu, H. (2016). Structural Basis and IP6 Requirement for Pds5-Dependent Cohesin Dynamics. Mol Cell 62, 248–259. 10.1016/j.molcel.2016.02.033.

20. Alomer, R.M., da Silva, E.M.L., Chen, J., Piekarz, K.M., McDonald, K., Sansam, C.G., Sansam, C.L., and Rankin, S. (2017). Esco1 and Esco2 regulate distinct cohesin functions during cell cycle progression. Proc Natl Acad Sci U S A 114, 9906–9911. 10.1073/pnas.1708291114.

21. Ben-Shahar, T.R., Heeger, S., Lehane, C., East, P., Flynn, H., Skehel, M., and Uhlmann, F. (2008). Eco1-dependent cohesin acetylation during establishment of sister chromatid cohesion. Science 321, 563–566. 321/5888/563 [pii] 10.1126/science.1157774 [doi].

22. Hou, F., and Zou, H. (2005). Two human orthologues of Eco1/Ctf7 acetyltransferases are both required for proper sister-chromatid cohesion. Mol Biol Cell 16, 3908–3918. 10.1091/mbc.e04-12-1063.

23. Rowland, B.D., Roig, M.B., Nishino, T., Kurze, A., Uluocak, P., Mishra, A., Beckouet, F., Underwood, P., Metson, J., Imre, R., et al. (2009). Building sister chromatid cohesion: smc3 acetylation counteracts an antiestablishment activity. Mol Cell 33, 763–774. S1097-2765(09)00144-0 [pii] 10.1016/j.molcel.2009.02.028 [doi].

24. Unal, E., Heidinger-Pauli, J.M., Kim, W., Guacci, V., Onn, I., Gygi, S.P., and Koshland, D.E. (2008). A molecular determinant for the establishment of sister chromatid cohesion. Science 321, 566–569. 321/5888/566 [pii] 10.1126/science.1157880 [doi].

25. Zhang, J., Shi, X., Li, Y., Kim, B.J., Jia, J., Huang, Z., Yang, T., Fu, X., Jung, S.Y., Wang, Y., et al. (2008). Acetylation of Smc3 by Eco1 is required for s phase sister chromatid cohesion in both human and yeast. Mol Cell 31, 143–151. S1097-2765(08)00420-6 [pii] 10.1016/j.molcel.2008.06.006 [doi].

26. Zheng, G., Kanchwala, M., Xing, C., and Yu, H. (2018). MCM2-7-dependent cohesin loading during S phase promotes sister-chromatid cohesion. Elife 7. 10.7554/eLife.33920.

27. Nishiyama, T., Ladurner, R., Schmitz, J., Kreidl, E., Schleiffer, A., Bhaskara, V., Bando, M., Shirahige, K., Hyman, A.A., Mechtler, K., and Peters, J.M. (2010). Sororin mediates sister chromatid cohesion by antagonizing Wapl. Cell 143, 737–749. 10.1016/j.cell.2010.10.031.

28. Rankin, S., Ayad, N.G., and Kirschner, M.W. (2005). Sororin, a substrate of the anaphase-promoting complex, is required for sister chromatid cohesion in vertebrates. Mol Cell 18, 185–200. S1097-2765(05)01187-1 [pii] 10.1016/j.molcel.2005.03.017 [doi].

29. Nishiyama, T., Sykora, M.M., Huis In ‘t Veld, P.J., Mechtler, K., and Peters, J.M. (2013). Aurora B and Cdk1 mediate Wapl activation and release of acetylated cohesin from chromosomes by phosphorylating Sororin. Proc Natl Acad Sci U S A 110, 13404–13409. 10.1073/pnas.1305020110.

30. Sumara, I., Vorlaufer, E., Stukenberg, P.T., Kelm, O., Redemann, N., Nigg, E.A., and Peters, J.M. (2002). The dissociation of cohesin from chromosomes in prophase is regulated by Pololike kinase. Mol Cell 9, 515–525.

31. Kitajima, T.S., Sakuno, T., Ishiguro, K.I., Iemura, S.I., Natsume, T., Kawashima, S.A., and Watanabe, Y. (2006). Shugoshin collaborates with protein phosphatase 2A to protect cohesin. Nature.

32. Liu, H., Rankin, S., and Yu, H. (2013). Phosphorylation-enabled binding of SGO1-PP2A to cohesin protects sororin and centromeric cohesion during mitosis. Nat Cell Biol 15, 40–49. 10.1038/ncb2637.

33. Liang, C., Chen, Q., Yi, Q., Zhang, M., Yan, H., Zhang, B., Zhou, L., Zhang, Z., Qi, F., Ye, S., and Wang, F. (2018). A kinase-dependent role for Haspin in antagonizing Wapl and protecting mitotic centromere cohesion. EMBO Rep 19, 43–56. 10.15252/embr.201744737.

34. Zhou, L., Liang, C., Chen, Q., Zhang, Z., Zhang, B., Yan, H., Qi, F., Zhang, M., Yi, Q., Guan, Y., et al. (2017). The N-Terminal Non-Kinase-Domain-Mediated Binding of Haspin to Pds5B Protects Centromeric Cohesion in Mitosis. Curr Biol 27, 992–1004. 10.1016/j.cub.2017.02.019.

35. London, N., and Biggins, S. (2014). Signalling dynamics in the spindle checkpoint response. Nat Rev Mol Cell Biol 15, 736–748. 10.1038/nrm3888.

36. Daum, J.R., Potapova, T.A., Sivakumar, S., Daniel, J.J., Flynn, J.N., Rankin, S., and Gorbsky, G.J. (2011). Cohesion Fatigue Induces Chromatid Separation in Cells Delayed at Metaphase. Curr Biol. 10.1016/j.cub.2011.05.032.

37. Stevens, D., Gassmann, R., Oegema, K., and Desai, A. (2011). Uncoordinated loss of chromatid cohesion is a common outcome of extended metaphase arrest. PLoS ONE 6, e22969. 10.1371/journal.pone.0022969.

38. Eot-Houllier, G., Fulcrand, G., Watanabe, Y., Magnaghi-Jaulin, L., and Jaulin, C. (2008). Histone deacetylase 3 is required for centromeric H3K4 deacetylation and sister chromatid cohesion. Genes Dev 22, 2639–2644. 22/19/2639 [pii] 10.1101/gad.484108.

39. Han, X., Piao, L., Zhuang, Q., Yuan, X., Liu, Z., and He, X. (2018). The role of histone lysine methyltransferase NSD3 in cancer. Onco Targets Ther 11, 3847–3852. 10.2147/ott.s166006.

40. Lucio-Eterovic, A.K., and Carpenter, P.B. (2011). An open and shut case for the role of NSD proteins as oncogenes. Transcription 2, 158–161. 10.4161/trns.2.4.16217.

41. Li, Y., Trojer, P., Xu, C.F., Cheung, P., Kuo, A., Drury, W.J., 3rd, Qiao, Q., Neubert, T.A., Xu, R.M., Gozani, O., and Reinberg, D. (2009). The target of the NSD family of histone lysine methyltransferases depends on the nature of the substrate. J Biol Chem 284, 34283–34295. 10.1074/jbc.M109.034462.

42. Wagner, E.J., and Carpenter, P.B. (2012). Understanding the language of Lys36 methylation at histone H3. Nat Rev Mol Cell Biol 13, 115–126. 10.1038/nrm3274.

43. Li, W., Tian, W., Yuan, G., Deng, P., Sengupta, D., Cheng, Z., Cao, Y., Ren, J., Qin, Y., Zhou, Y., et al. (2021). Molecular basis of nucleosomal H3K36 methylation by NSD methyltransferases. Nature 590, 498–503. 10.1038/s41586-020-03069-8.

44. Rahman, S., Sowa, M.E., Ottinger, M., Smith, J.A., Shi, Y., Harper, J.W., and Howley, P.M. (2011). The Brd4 extraterminal domain confers transcription activation independent of pTEFb by recruiting multiple proteins, including NSD3. Mol Cell Biol 31, 2641–2652. 10.1128/mcb.01341-10.

45. Yuan, G., Flores, N.M., Hausmann, S., Lofgren, S.M., Kharchenko, V., Angulo-Ibanez, M., Sengupta, D., Lu, X., Czaban, I., Azhibek, D., et al. (2021). Elevated NSD3 histone methylation activity drives squamous cell lung cancer. Nature 590, 504–508. 10.1038/s41586-020-03170-y.

46. Linares-Saldana, R., Kim, W., Bolar, N.A., Zhang, H., Koch-Bojalad, B.A., Yoon, S., Shah, P.P., Karnay, A., Park, D.S., Luppino, J.M., et al. (2021). BRD4 orchestrates genome folding to promote neural crest differentiation. Nat Genet 53, 1480–1492. 10.1038/s41588-021-00934-8.

47. Angrand, P.O., Apiou, F., Stewart, A.F., Dutrillaux, B., Losson, R., and Chambon, P. (2001). NSD3, a new SET domain-containing gene, maps to 8p12 and is amplified in human breast cancer cell lines. Genomics 74, 79–88. 10.1006/geno.2001.6524.

48. Kim, S.M., Kee, H.J., Eom, G.H., Choe, N.W., Kim, J.Y., Kim, Y.S., Kim, S.K., Kook, H., Kook, H., and Seo, S.B. (2006). Characterization of a novel WHSC1-associated SET domain protein with H3K4 and H3K27 methyltransferase activity. Biochem Biophys Res Commun 345, 318–323. 10.1016/j.bbrc.2006.04.095.

49. Bennett, R.L., Swaroop, A., Troche, C., and Licht, J.D. (2017). The Role of Nuclear Receptor-Binding SET Domain Family Histone Lysine Methyltransferases in Cancer. Cold Spring Harb Perspect Med 7. 10.1101/cshperspect.a026708.

50. Zhang, M., Yang, Y., Zhou, M., Dong, A., Yan, X., Loppnau, P., Min, J., and Liu, Y. (2021). Histone and DNA binding ability studies of the NSD subfamily of PWWP domains. Biochem Biophys Res Commun 569, 199–206. 10.1016/j.bbrc.2021.07.017.

51. Wu, H., Zeng, H., Lam, R., Tempel, W., Amaya, M.F., Xu, C., Dombrovski, L., Qiu, W., Wang, Y., and Min, J. (2011). Structural and histone binding ability characterizations of human PWWP domains. PLoS One 6, e18919. 10.1371/journal.pone.0018919.

52. Vermeulen, M., Eberl, H.C., Matarese, F., Marks, H., Denissov, S., Butter, F., Lee, K.K., Olsen, J.V., Hyman, A.A., Stunnenberg, H.G., and Mann, M. (2010). Quantitative interaction proteomics and genome-wide profiling of epigenetic histone marks and their readers. Cell 142, 967–980. 10.1016/j.cell.2010.08.020.

53. Zhou, Z., Thomsen, R., Kahns, S., and Nielsen, A.L. (2010). The NSD3L histone methyltransferase regulates cell cycle and cell invasion in breast cancer cells. Biochem Biophys Res Commun 398, 565–570. 10.1016/j.bbrc.2010.06.119.

54. Shen, C., Ipsaro, J.J., Shi, J., Milazzo, J.P., Wang, E., Roe, J.S., Suzuki, Y., Pappin, D.J., Joshua-Tor, L., and Vakoc, C.R. (2015). NSD3-Short Is an Adaptor Protein that Couples BRD4 to the CHD8 Chromatin Remodeler. Mol Cell 60, 847–859. 10.1016/j.molcel.2015.10.033.

55. Liu, H., Jia, L., and Yu, H. (2013). Phospho-H2A and Cohesin Specify Distinct Tension-Regulated Sgo1 Pools at Kinetochores and Inner Centromeres. Curr Biol. 10.1016/j.cub.2013.07.078.

56. Zhou, L., Liang, C., Chen, Q., Zhang, Z., Zhang, B., Yan, H., Qi, F., Zhang, M., Yi, Q., Guan, Y., et al. (2017). The N-Terminal Non-Kinase-Domain-Mediated Binding of Haspin to Pds5B Protects Centromeric Cohesion in Mitosis. Curr Biol 27, 992–1004. 10.1016/j.cub.2017.02.019.

57. Goto, Y., Yamagishi, Y., Shintomi-Kawamura, M., Abe, M., Tanno, Y., and Watanabe, Y. (2017). Pds5 Regulates Sister-Chromatid Cohesion and Chromosome Bi-orientation through a Conserved Protein Interaction Module. Current Biology 27, 1005–1012. 10.1016/j.cub.2017.02.066.

58. Li, W., Tian, W., Yuan, G., Deng, P., Sengupta, D., Cheng, Z., Cao, Y., Ren, J., Qin, Y., Zhou, Y., et al. (2021). Molecular basis of nucleosomal H3K36 methylation by NSD methyltransferases. Nature 590, 498–503. 10.1038/s41586-020-03069-8.

59. Hahn, M., Dambacher, S., Dulev, S., Kuznetsova, A.Y., Eck, S., Worz, S., Sadic, D., Schulte, M., Mallm, J.P., Maiser, A., et al. (2013). Suv4-20h2 mediates chromatin compaction and is important for cohesin recruitment to heterochromatin. Genes Dev 27, 859–872. 10.1101/gad.210377.112.

60. Ritchie, K., Seah, C., Moulin, J., Isaac, C., Dick, F., and Berube, N.G. (2008). Loss of ATRX leads to chromosome cohesion and congression defects. J Cell Biol 180, 315–324. jcb.200706083 [pii] 10.1083/jcb.200706083 [doi].

61. Shen, C., Ipsaro, J.J., Shi, J., Milazzo, J.P., Wang, E., Roe, J.S., Suzuki, Y., Pappin, D.J., Joshua-Tor, L., and Vakoc, C.R. (2015). NSD3-Short Is an Adaptor Protein that Couples BRD4 to the CHD8 Chromatin Remodeler. Mol Cell 60, 847–859. 10.1016/j.molcel.2015.10.033.

62. Heidari, N., Phanstiel, D.H., He, C., Grubert, F., Jahanbani, F., Kasowski, M., Zhang, M.Q., and Snyder, M.P. (2014). Genome-wide map of regulatory interactions in the human genome. Genome Res 24, 1905–1917. 10.1101/gr.176586.114.

63. DeMare, L.E., Leng, J., Cotney, J., Reilly, S.K., Yin, J., Sarro, R., and Noonan, J.P. (2013). The genomic landscape of cohesin-associated chromatin interactions. Genome Res 23, 1224–1234. 10.1101/gr.156570.113.

64. Kagey, M.H., Newman, J.J., Bilodeau, S., Zhan, Y., Orlando, D.A., van Berkum, N.L., Ebmeier, C.C., Goossens, J., Rahl, P.B., Levine, S.S., et al. (2010). Mediator and cohesin connect gene expression and chromatin architecture. Nature 467, 430–435. 10.1038/nature09380.

65. Pherson, M., Misulovin, Z., Gause, M., and Dorsett, D. (2019). Cohesin occupancy and composition at enhancers and promoters are linked to DNA replication origin proximity in. Genome Res 29, 602–612. 10.1101/gr.243832.118.

66. Linares-Saldana, R., Kim, W., Bolar, N.A., Zhang, H., Koch-Bojalad, B.A., Yoon, S., Shah, P.P., Karnay, A., Park, D.S., Luppino, J.M., et al. (2021). BRD4 orchestrates genome folding to promote neural crest differentiation. Nat Genet 53, 1480–1492. 10.1038/s41588-021-00934-8.

67. Lopez-Serra, L., Kelly, G., Patel, H., Stewart, A., and Uhlmann, F. (2014). The Scc2-Scc4 complex acts in sister chromatid cohesion and transcriptional regulation by maintaining nucleosome-free regions. Nat Genet 46, 1147–1151. 10.1038/ng.3080.

68. Muñoz, S., Minamino, M., Casas-Delucchi, C.S., Patel, H., and Uhlmann, F. (2019). A Role for Chromatin Remodeling in Cohesin Loading onto Chromosomes. Mol Cell 74, 664–673 e665. 10.1016/j.molcel.2019.02.027.

69. Muñoz, S., Passarelli, F., and Uhlmann, F. (2020). Conserved roles of chromatin remodellers in cohesin loading onto chromatin. Curr Genet 66, 951–956. 10.1007/s00294-020-01075-x.

70. Muñoz, S., Jones, A., Bouchoux, C., Gilmore, T., Patel, H., and Uhlmann, F. (2022). Functional crosstalk between the cohesin loader and chromatin remodelers. Nat Commun 13, 7698. 10.1038/s41467-022-35444-6.

71. Hakimi, M.A., Bochar, D.A., Schmiesing, J.A., Dong, Y., Barak, O.G., Speicher, D.W., Yokomori, K., and Shiekhattar, R. (2002). A chromatin remodelling complex that loads cohesin onto human chromosomes. Nature 418, 994–998.

72. Eid, R., Demattei, M.V., Episkopou, H., Augé-Gouillou, C., Decottignies, A., Grandin, N., and Charbonneau, M. (2015). Genetic Inactivation of ATRX Leads to a Decrease in the Amount of Telomeric Cohesin and Level of Telomere Transcription in Human Glioma Cells. Mol Cell Biol 35, 2818–2830. 10.1128/mcb.01317-14.

73. Kernohan, K.D., Jiang, Y., Tremblay, D.C., Bonvissuto, A.C., Eubanks, J.H., Mann, M.R., and Bérubé, N.G. (2010). ATRX partners with cohesin and MeCP2 and contributes to developmental silencing of imprinted genes in the brain. Dev Cell 18, 191–202. 10.1016/j.devcel.2009.12.017.

74. Zuin, J., Franke, V., van Ijcken, W.F., van der Sloot, A., Krantz, I.D., van der Reijden, M.I., Nakato, R., Lenhard, B., and Wendt, K.S. (2014). A cohesin-independent role for NIPBL at promoters provides insights in CdLS. PLoS Genet 10, e1004153. 10.1371/journal.pgen.1004153.

75. Kuo, A.J., Cheung, P., Chen, K., Zee, B.M., Kioi, M., Lauring, J., Xi, Y., Park, B.H., Shi, X., Garcia, B.A., et al. (2011). NSD2 links dimethylation of histone H3 at lysine 36 to oncogenic programming. Mol Cell 44, 609–620. 10.1016/j.molcel.2011.08.042.

76. Zhu, L., Li, Q., Wong, S.H., Huang, M., Klein, B.J., Shen, J., Ikenouye, L., Onishi, M., Schneidawind, D., Buechele, C., et al. (2016). ASH1L Links Histone H3 Lysine 36 Dimethylation to MLL Leukemia. Cancer Discov 6, 770–783. 10.1158/2159-8290.cd-16-0058.

77. Sumara, I., Vorlaufer, E., Gieffers, C., Peters, B.H., and Peters, J.M. (2000). Characterization of vertebrate cohesin complexes and their regulation in prophase. J Cell Biol 151, 749–762. 10.1083/jcb.151.4.749.

78. Nishiyama, T., Ladurner, R., Schmitz, J., Kreidl, E., Schleiffer, A., Bhaskara, V., Bando, M., Shirahige, K., Hyman, A.A., Mechtler, K., and Peters, J.M. (2010). Sororin mediates sister chromatid cohesion by antagonizing Wapl. Cell 143, 737–749. 10.1016/j.cell.2010.10.031.

79. Watrin, E., and Legagneux, V. (2005). Contribution of hCAP-D2, a non-SMC subunit of condensin I, to chromosome and chromosomal protein dynamics during mitosis. Mol Cell Biol 25, 740–750. 10.1128/mcb.25.2.740-750.2005.

80. Kitajima, T.S., Hauf, S., Ohsugi, M., Yamamoto, T., and Watanabe, Y. (2005). Human Bub1 defines the persistent cohesion site along the mitotic chromosome by affecting Shugoshin localization. Curr Biol 15, 353–359.

81. Schindelin, J., Arganda-Carreras, I., Frise, E., Kaynig, V., Longair, M., Pietzsch, T., Preibisch, S., Rueden, C., Saalfeld, S., Schmid, B., et al. (2012). Fiji: an open-source platform for biological-image analysis. Nat Methods 9, 676–682. 10.1038/nmeth.2019.

82. Schmitz, J., Watrin, E., Lenart, P., Mechtler, K., and Peters, J.M. (2007). Sororin Is Required for Stable Binding of Cohesin to Chromatin and for Sister Chromatid Cohesion in Interphase. Curr Biol.

