## Supplemental material for "The histone methyltransferase NSD3 contributes to sister chromatid cohesion and to cohesin loading at mitotic exit"

### Supplementary material

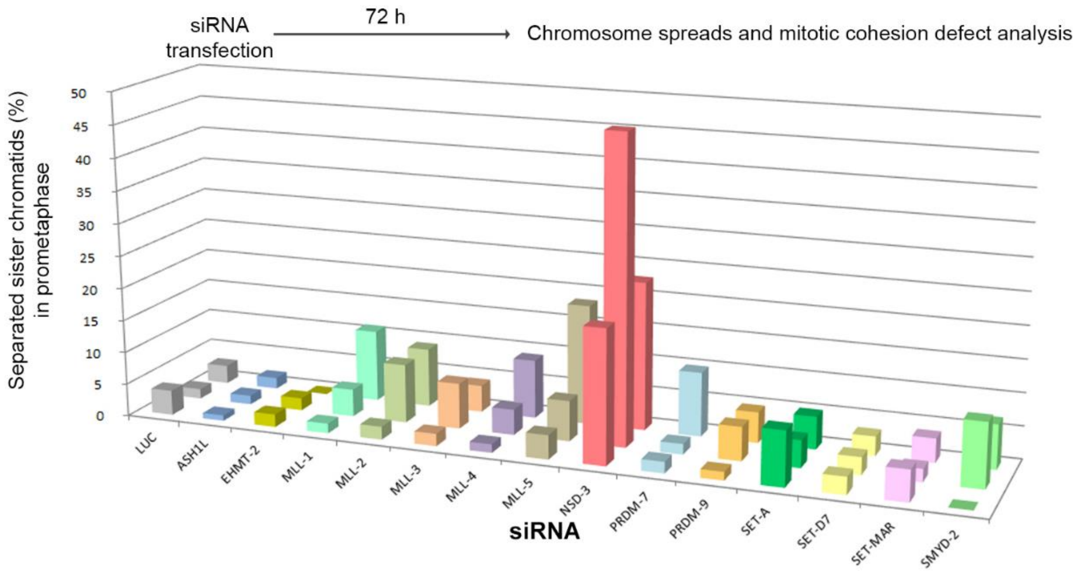


**Figure S1: Screening for methyltransferases that contain a SET domain and are involved in preventing sister chromatid separation during mitosis**

**Figure S1: Screening for methyltransferases that contain a SET domain and are involved in preventing sister chromatid separation during mitosis**. For each of the 14 evaluated methyltransferases, 3 different siRNAs were tested. Cells were transfected for 72 h before fixation and chromosome spreading. Between 300 and 500 prometaphase cells were counted, and the proportion of mitotic cells with separated chromatids was assessed according to the phenotypes shown in Figure 1A.

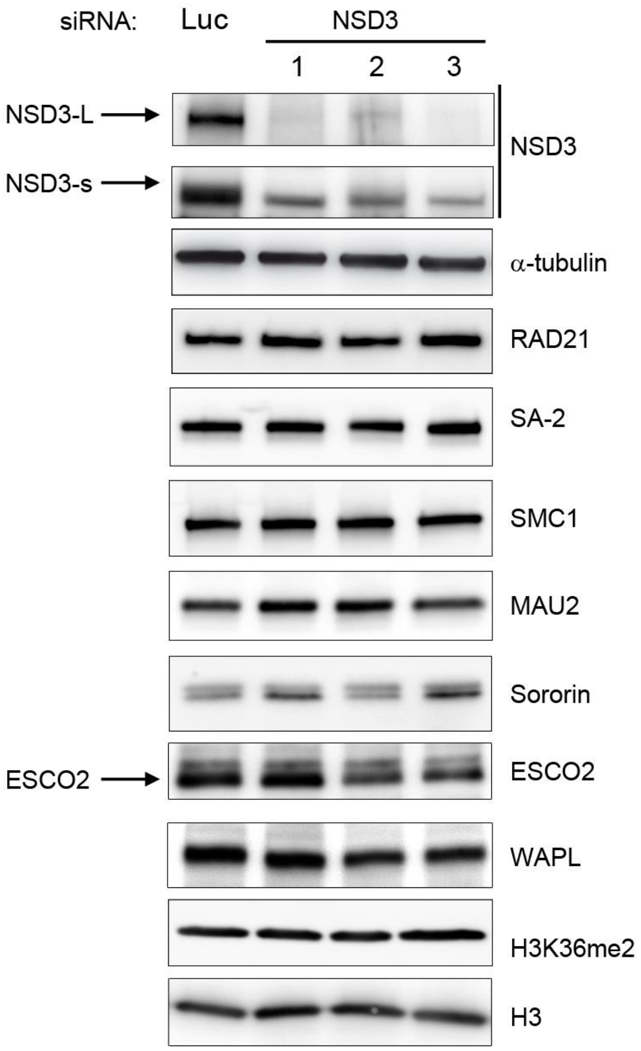


**Figure S2: NSD3 depletion does not affect expression levels of the core cohesin complex components or associated regulatory partners, nor does it affect the global levels of H3K36 di-methylation**

**Figure S2: NSD3 depletion does not affect expression levels of the core cohesin complex components or associated regulatory partners, nor does it affect the global levels of H3K36 di-methylation**. Whole-cell extracts from cells transfected for 72 h with three different NSD3 siRNAs were prepared and analysed by western blot. α-tubulin and histone H3 were used as the cytoplasmic and nuclear loading markers, respectively. Note the presence of a non-specific band above the ESCO2 band.

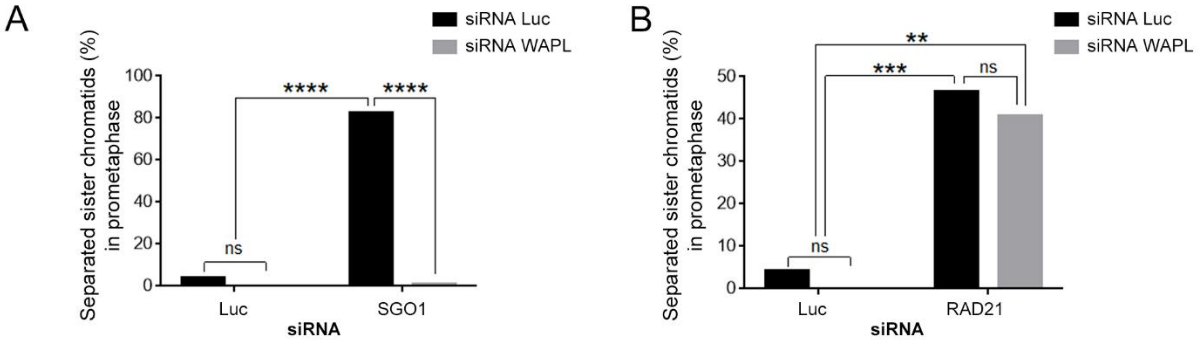


Figure S3: WAPL prevents induction of mitotic cohesion defects when RAD21 is depleted but not when SGO1 is depleted

**Figure S3: WAPL prevents induction of mitotic cohesion defects when RAD21 is depleted but not when SGO1 is depleted**. HeLa cells were transfected with Luc, SGO1, or RAD21 siRNA. After 24 h, a second transfection was performed with Luc or WAPL siRNA. After further incubation for 24 h, the cells were harvested, and the chromosomes were spread and labelled with DAPI. The proportion of mitotic cells with separated chromatids is shown as compared to the total mitotic cells. ≥ 150 prometaphase cells were counted for each condition. ns: not significant, **: $P \leq 0.01$, ***: $P \leq 0.001$, ****: $P \leq 0.0001$ (Chi-square test).

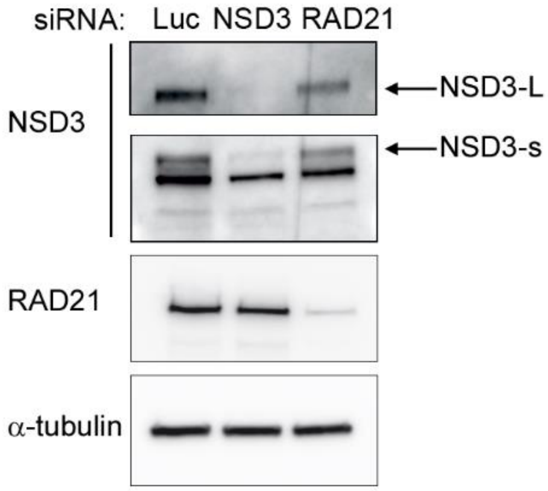


**Figure S4: Efficiency of the depletion of NSD3 and RAD21 in cells used for DNA FISH experiments**

**Figure S4: Efficiency of the depletion of NSD3 and RAD21 in cells used for DNA FISH experiments**. HeLa cells were transfected for 8 h with siRNA. They were then subjected to a double thymidine-block followed by a 6-h release to enrich G2 cells. For each condition, a fraction of the cell population was then used to prepare whole-cell extracts for western blot analysis in order to identify any protein extinctions caused by siRNA.

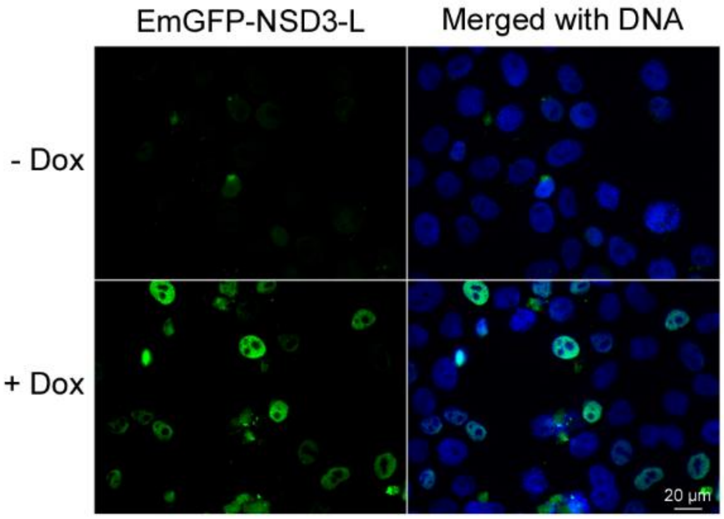


**Figure S5: HeLa cell line expressing an inducible NSD3 protein fused to a EmGFP tag**

**Figure S5: HeLa cell line expressing an inducible NSD3 protein fused to a EmGFP tag**. Cells were left alone or treated with doxycycline for 48 h, fixed, then labelled with blue DAPI. Cells expressing EmGFP-NSD3-L were observable in green. Shown are representative images of cells with or without treatment with doxycycline. Note that not all cells were induced, despite clonal selection after doxycyline treatment.

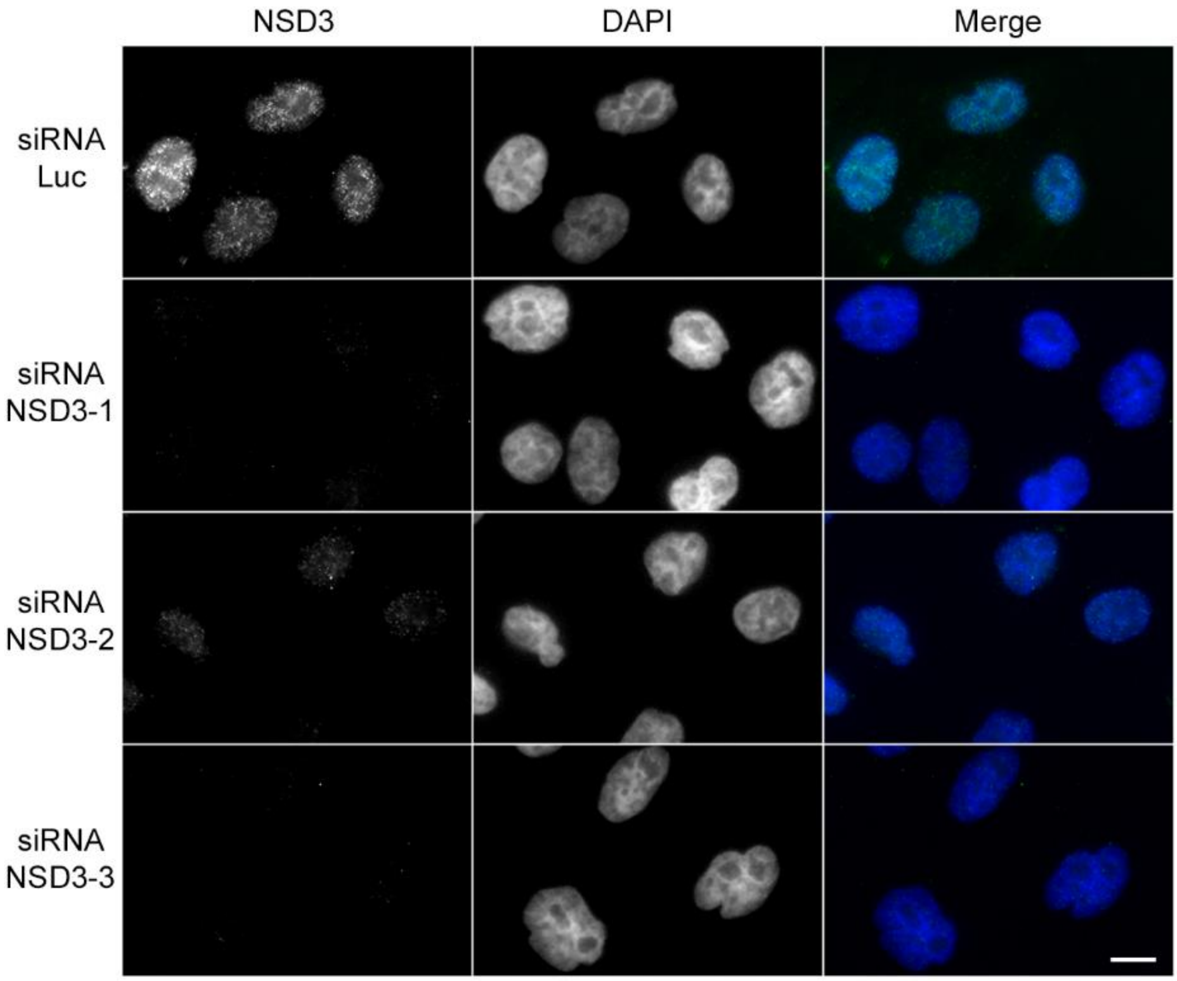


**Figure S6: Validation of the NSD3 antibody for immunostaining**

**Figure S6: Validation of the NSD3 antibody for immunostaining**. HeLa cells were transfected with the indicated siRNAs for 48 h. The soluble cell content was then extracted, subjected to immunofluorescence with anti-NSD3 antibodies, and stained with DAPI. Scale bar: 10 µm.

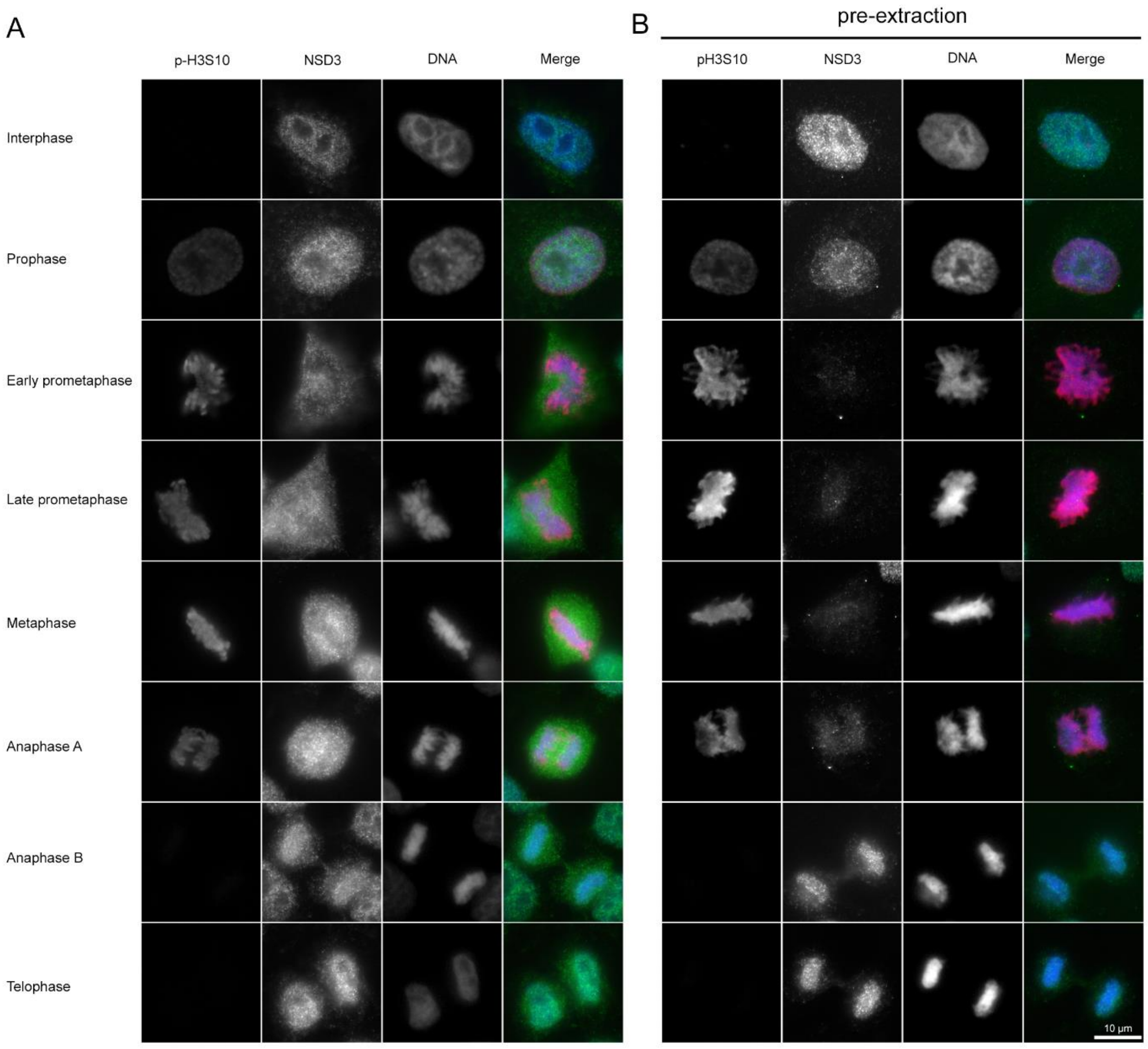


Figure S7: NSD3 localisation during the cell cycle

**Figure S7: NSD3 localisation during the cell cycle**. Proliferative HeLa cells were fixed immediately (A) or 1 min after incubation with 0.1% triton to extract the soluble cell content (B). Cells were then immunolabeled with anti-NSD3 antibodies. Anti-phosphorylated H3S10 antibodies were used as a marker for the prophase through early anaphase stages of cell division.

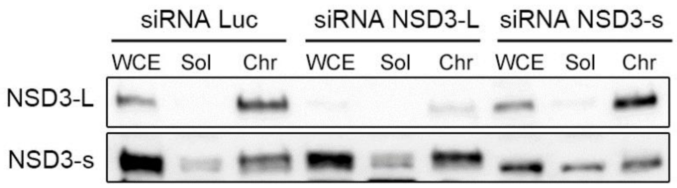


**Figure S8: Depletion of NSD3 isoforms for kollerin- and cohesin-loading experiments**

**Figure S8: Depletion of NSD3 isoforms for kollerin- and cohesin-loading experiments.** WCE, Sol and Chr correspond to whole-cell extracts and soluble and chromatin fractions, respectively (see Materials and methods). In contrast to Figure 5D, western blots against NSD3 isoforms are represented here on the same line to show that NSD3-s was correctly depleted. Note that after the specific depletion of NSD3-s, a non-specific band was still present on the gel.

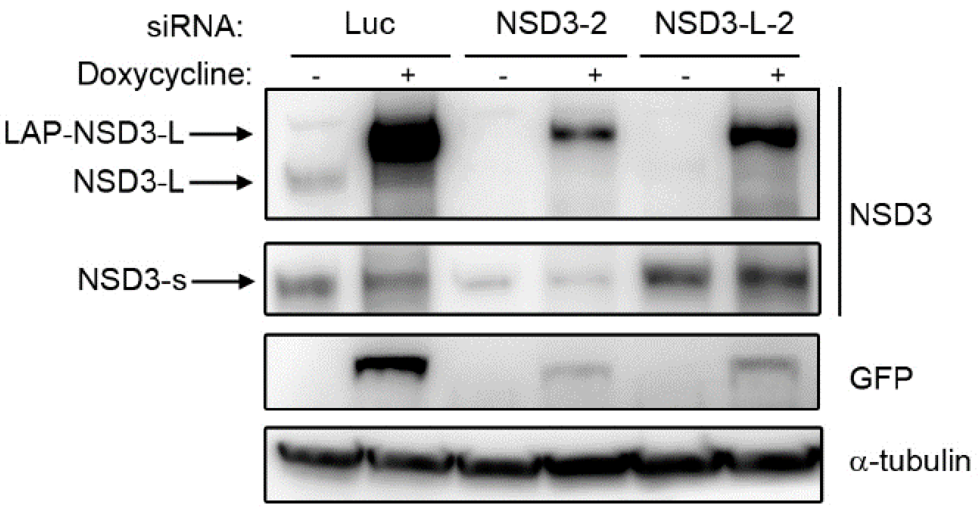


**Figure S9: Expression levels of endogenous and exogenous NSD3 in rescue experiments**

**Figure S9: Expression levels of endogenous and exogenous NSD3 in rescue experiments**. Western blot analysis of exogenous and endogenous forms of NSD3 following endogenous NSD3 depletion and concomitant induction of wild-type EmGFP-NSD3-L expression. The band corresponding to the EmGFP-tag fused with NSD3-L, detected only after doxycycline induction, migrated more slowly than those of endogenous NSD3-L.

**Supplementary Movie 1: Time-lapse images of H2B-mCherry expression in a HeLa cell.**

**Supplementary Movie 2: Time-lapse images of EmGFP-NSD3-L expression in a HeLa cell.**

**Supplementary Movie 3: Merged time-lapse images of H2B-mCherry and EmGFP-NSD3-L expression in a HeLa cell.**
